# DeepTRACE: Flexible Machine Learning for Analysis and Discovery in Single Molecule Tracks

**DOI:** 10.1101/2025.05.15.654348

**Authors:** Oliver J Pambos, Jacob AR Wright, Achillefs N Kapanidis

## Abstract

DeepTRACE is a machine learning tool for analysing long, complex single-molecule tracks in living cells; learning from sequences of molecular events using spatial, temporal, and photometric context. It traces how molecular processes unfold over time and space, incorporating past interactions, subcellular location, and photometric properties. DeepTRACE requires only a few hundred annotated tracks and minutes of CPU training, yet outperforms traditional methods and supports extensive downstream analysis, including the discovery of relationships absent from the training data.

## Main Text

Single-Molecule Localisation Microscopy (SMLM) has provided stunning insight into biological processes, with single-molecule tracking representing one of its most powerful modes. In a typical tracking experiment, the chronological sequence of a molecule’s positions is connected into a track, revealing its motion. Such experiments have provided insight into fundamental biological processes, such as target search on the chromosome [1] and Z-ring organisation during bacterial cell division [2].

Early tracking experiments were limited to the millisecond-to-second timescale due to fast photobleaching of the available fluorescent proteins; established analysis pipelines were thus designed for shorter, non-transitioning tracks, which provided mostly truncated snapshots of biological processes and did not report directly on dynamics.

Recently, single-molecule tracks have been extended to the multi-minute timescale, due to the introduction of brighter and more stable probes [3–8], new labelling strategies [9, 10], replenishing probes [11, 12], and temporally-patterned excitation (stroboscopy, timelapse imaging). Such long-lived tracks hold tremendous promise for capturing entire biological processes (e.g. transcription, translation, cell division) that unfold over minutes, catalysing new discoveries.

This potential has remained unrealised, largely because current track analysis involves rigid constraints, such as processing only pure diffusive states [13,14], truncated snapshots of tracks [15], memoryless transitions [16–18], fixed classification thresholds [15, 19], rigidly-defined theoretical diffusion classes [20], fixed numbers of classes [21], and a lack of spatial context [13, 15–21]. Biological processes, however, follow complex sequences of events, with actions depending upon past interactions, current molecular conformation, and the cellular location at which they occur.

To address these limitations, we developed DeepTRACE (**D**eep learning-based **T**rack **R**ecognition, **A**nalysis, and **C**lassification of **E**vents), a flexible, open-source, cross-platform, GUI-driven software for single-molecule track segmentation and analysis. DeepTRACE combines temporal and spatial context with photometric information and user-supplied natural language labels to train bespoke machine learning (ML) models, using an interface that requires no coding or ML experience.

DeepTRACE supports two separate modes for generating training labels. First, users can train directly on simulated single-molecule tracking data, using a simple tabular format containing localisation-level class labels as ground truth. Alternatively, users can directly annotate real experimental data using DeepTRACE’s interface. This interactive tool presents the video of each molecule alongside its 2D track and a synchronised feature time series (e.g. *step size* vs time), allowing intuitive segmentation of molecular behaviour. While manual annotation inevitably reflects human judgement, using experimental perturbations (e.g. antibiotics, protein over-expression) to enhance rare or hard-to-identify states both improves the reliability of human-generated labels and increases the density of key states, thereby enabling more effective evaluation in the native, unperturbed system — a strategy analogous to Gamma Mixture Model (γMM) methods, where perturbations are used to fix model states for subsequent application to unperturbed conditions.

DeepTRACE combines tracking data with segmented cell boundaries to perform feature engineering; wherein powerful features (e.g. *path straightness* or *distance to cell membrane*) which encode higher order spatio-temporal or photometric properties, are constructed from raw features such as particle coordinates (steps 1-2, **Fig. 1a**). Tracks are annotated with natural language labels (step 3, **Fig. 1a**) using manual annotation, simulations, or both. The user then selects features likely to reflect the biological process studied, and these train a custom ML model (steps 4-5, **Fig. 1a**). Finally, DeepTRACE segments unseen tracks into subtracks using sequence-to-sequence classification with a sliding window voting approach (**Fig. 1b, Online Methods**) enabling robust, rapid segmentation of tracks of any length for downstream analysis (e.g. diffusion analysis, spatial mapping, kinetics).

**Figure 1.**
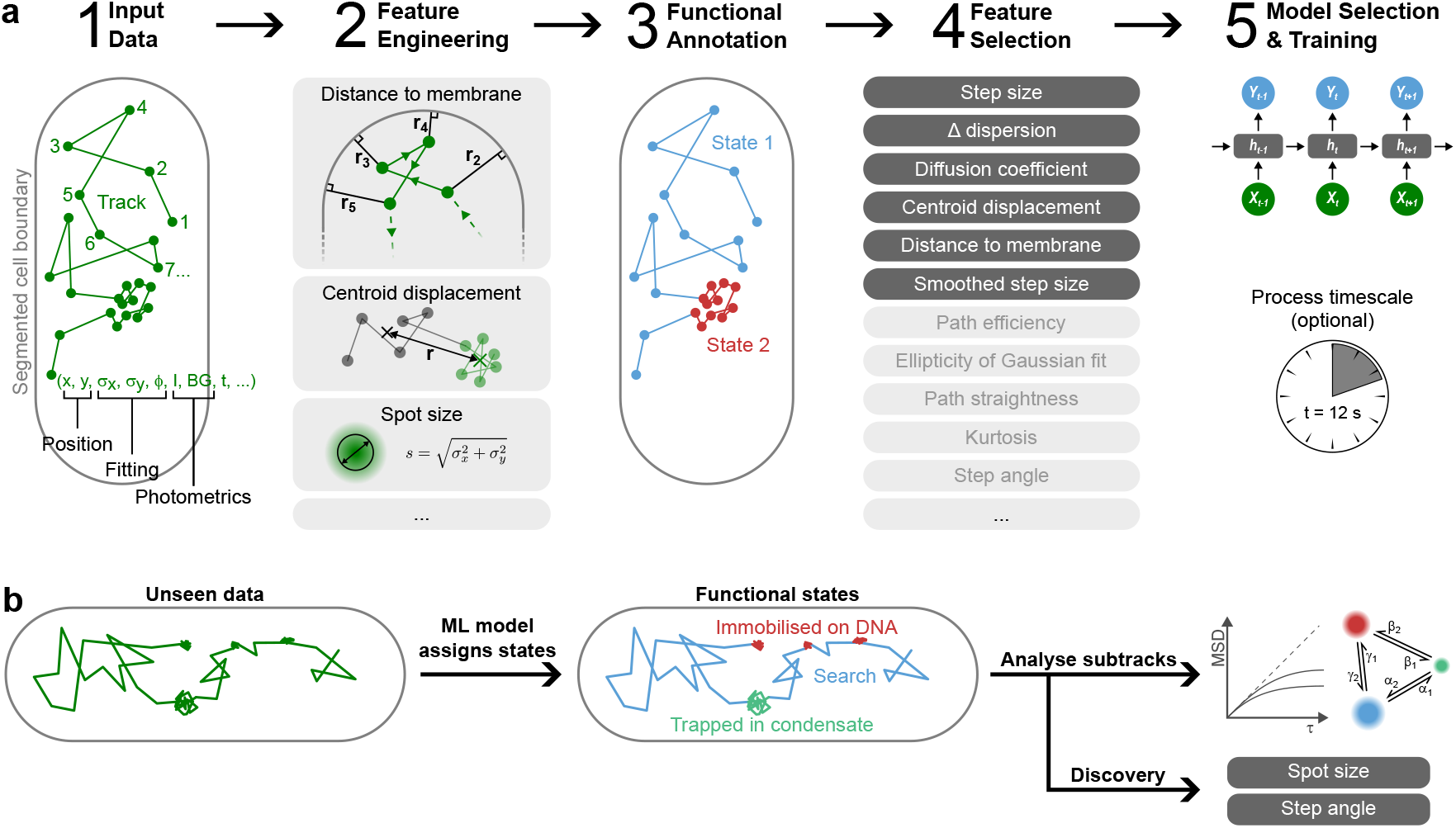
The DeepTRACE workflow. **a**, Raw tracking, photometric, and cell segmentation data from a small test set are used to engineer a large number of features encoding different aspects of each molecule’s behaviour. Biological states are annotated, either through human annotation of real experimental data, an objective ground truth from theoretical simulations, or a combination of the two. The user then identifies features likely to be predictive of the biological process, and these features are used to train a bespoke machine learning model. **b**, The trained model segments unseen tracks into subtracks representing biological states, which are used in downstream analysis. Finally, the system identifies any unseen relationships between the data and features not used by the model, for hypothesis-free discovery.

As feature selection is crucial, DeepTRACE’s GUI offers extensive visual guidance to select powerful non-redundant features (**Extended Data Fig. 1-2**).

We evaluated DeepTRACE using both simulated and real experimental data. To provide an unambiguous ground truth, we first simulated realistic particle motions inside a 3D model cell (**Fig. 2a**) using Smoldyn [22]. As a model cell geometry, we chose a rod-shaped bacterial cell; this tight spatial confinement masks the anomalous diffusion exponent that otherwise simplifies the task of motion identification in larger eukaryotic cells or extended membranes (**Extended Data Fig. 3**). We used these 3D tracks to construct a series of fluorescence video frames using SMeagol [23], producing videos that incorporate key experimental parameters (e.g. realistic point spread functions, diffusion and binding behaviours, diffusive transitions within frames, motion blur, camera noise, photoactivation, photobleaching, and confinement). We then processed the raw videos using a typical tracking pipeline [15,24], thus introducing directly the sources of noise in real particle-tracking applications, and imported the tracking data and ground truth into DeepTRACE.

**Figure 2.**
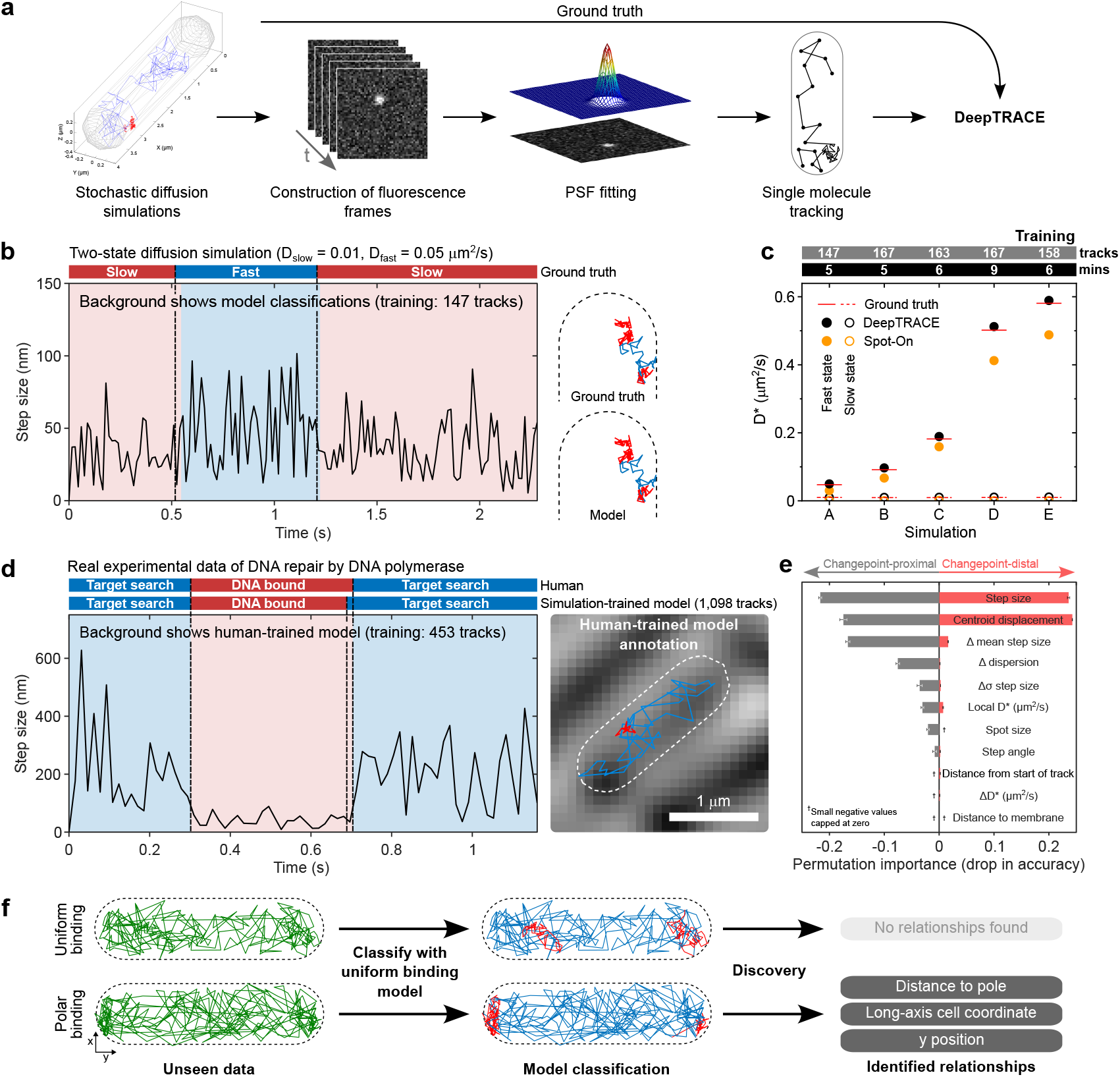
DeepTRACE analysis of simulated and real experimental data. **a**, The simulation pipeline; particle simulations with Smoldyn were rendered into fluorescence video frames using SMeagol, and processed using a typical tracking pipeline. **b**, Time series of simulated reversible switching between two diffusive states (*D* = 0.01 and 0.05 *μ*m^2^*/*s), showing classifications from a model trained on 147 tracks (inset colours), and ground truth (bars above). **c**, Performance comparison between DeepTRACE (black circles), and Spot-On [21] (orange circles) for inference of diffusion coefficients across five two-state diffusion simulations, compared to objective ground truth (red lines), using models trained with fewer than 200 tracks and 10 minutes of processing on a single CPU core (bars above). **d**, Time series of real Pol1 single-molecule tracks in live *E*.*coli* [24] showing classifications from a simulation-trained model and direct human annotation (bars above), compared to classifications from a human-trained model (inset colour, plotted track). **e**, Permutation importance showing the relative influence of features on classification of Pol1 tracks using a surrogate model. **f**, DeepTRACE’s *Discovery* tool evaluating simulated tracks of binding to a polar-localised plasmid using a model trained on uniformly-distributed binding events; identifying the existence of the new relationship with pole proximity absent from training data.

To test the versatility of DeepTRACE, we simulated a diverse range of behaviours: reversible transition between diffusive states (**Fig. 2b**), irreversible transport across the inner membrane (**Extended Data Fig. 4**), and reversible binding to a polar-localised plasmid (**Extended Data Fig. 5**). We performed each simulation with timelapse imaging (10 ms excitation, 190 ms dark intervals). We also simulated continuous recording in the two-state diffusion simulation with 15 ms exposures, common in trackingPALM [15, 24].

Reversible two-state diffusion simulations were segmented using a DeepTRACE model incorporating a Bidirectional Long Short-Term Memory network (BiLSTM) with a self-attention mechanism; shown in **Fig. 2b** as a time series of the *step size* feature indicating strong agreement with ground truth labels. We quantify its performance through state classification metrics and inference of simulated kinetic parameters, and compare with the established methods of γMM diffusion analysis [15], and the widely-used software Spot-On [21], using the objective ground truth across a wide range of diffusion coefficients (red lines, **Fig. 2c**). DeepTRACE outperformed both Spot-On and γMM analysis for inference of diffusion coefficients and state occupancies across all simulations (black and orange circles respectively, **Fig. 2c**; and teal circles, **Extended Data Fig. 6c**). This superior performance remained true even with DeepTRACE restricted to small training datasets (160 ± 8 tracks), small window size (25 frames), and rapid training (5-10 min) — all achieved without specialised ML hardware (laptop, single CPU core); see bars above plot **Fig. 2c**. By varying the dataset size and separation of simulated mobilities we found that DeepTRACE is capable of accurate segmentation using datasets equivalent in size to a single field-of-view recording, and with diffusion coefficients of common DNA-binding proteins (**Extended Data Fig. 7**). DeepTRACE also enables extensive downstream analysis and visualisation of class-specific properties; for example the dwell-time distributions from model-identified transport events (**Extended Data Fig. 4c**) and spatial mapping of all localisations classified as undergoing transport (showing bias towards the periphery; figure inset).

To assess DeepTRACE’s ability to learn directly from human annotation, we generated two-state diffusion simulations using experimentally-established diffusion coefficients (*D*_repair_ = 0.01 *μ*m^2^*/*s, *D*_search_ = 0.64 *μ*m^2^*/*s) for DNA polymerase I (Pol1) under 15 ms continuous imaging [24]. 197 tracks were manually annotated using the built-in *Human Annotation* tool and used to train a DeepTRACE model (167 training, 30 validation tracks). The trained model was then used to segment 1,292 tracks from a completely independent simulation, and DeepTRACE inferred kinetic parameters from the resulting subtracks. The same 1,292 tracks were analysed by Spot-On and γMM. Across all kinetic parameters, DeepTRACE returned values within 0.6 − 7% of ground truth, outperforming both γMM and Spot-On (**Extended Data Table 1**), demonstrating the ability to infer kinetic parameters with realistic protein mobilities using models trained exclusively on human annotation.

We then extended this approach to real experimental data, with the human annotation of 453 tracks from previously published experiments [24] of Pol1 repairing H_2_O_2_-induced DNA damage; a process involving ~ 2-second transient immobilisation events [19]. These labels were used to train a model, which subsequently annotated unseen data from independent experiments, showing strong agreement with separate expert human annotations of the same dataset (**Fig. 2d**). We also used the same model trained on human-annotation of real tracks to segment and analyse the simulated data, for which a ground truth exists. Remarkably, despite differences between experiment and the synthetic training set — e.g. reversed class balance, different localisation error, free diffusion of the simulated slow state, different binding times — the model again outperformed analysis of the same dataset using established methods (**Extended Data Table 1**).

To discover any unexpected relationships, DeepTRACE’s *Discovery* tool evaluates mutual information between classified states and features not used by the model, enabling hypothesis-free discovery. To illustrate this, we used simulations of a protein reversibly binding to a slowly diffusing plasmid located within the cell end-caps, resulting in it transiently adopting the plasmid’s mobility and spatial profile (**Extended Data Fig. 5**). By evaluating with a model trained on binding events distributed uniformly across the intracellular space, the *Discovery* tool correctly identified a relationship between pole proximity and diffusive state despite this relationship being absent from the model’s training data (**Fig. 2f**). No such relationships were identified in control data where events were uniformly distributed.

DeepTRACE’s efficient learning from small datasets arises mostly from the compression of key information into powerful engineered features (**Extended Data Fig. 7a** inset), strong regularisation, and track subsampling (which reframes each datapoint in different temporal contexts, capitalising on limited data; **Online Methods**). This segmentation enables improved sampling in mean-squared displacement (MSD) calculations, removes track truncation and track-length biases, and better separates diffusive states (**Extended Data Fig. 8**). DeepTRACE then performs diffusion calculations directly on segmented MSD data — in contrast to the standard approach of de-mixing populations from an ensemble distribution — achieving higher accuracy with fewer tracks. Consequently, a single field of view is sufficient to infer global properties (D*, transition rates, occupancies; **Extended Data Fig. 6, Extended Data Table 1**) which typically require ~ 80,000 tracks using standard track analysis [24].

The ability to further enhance changepoint detection through experimental perturbations was demonstrated by increasing class transition rates in simulations of two-state reversible diffusion simulations, and evaluating on separate unperturbed data, showing improvement across a wide range of training set sizes (**Extended Data Fig. 9a**).

We have restricted our comparisons to diffusion analysis as existing pipelines are limited to diffusion tasks; however DeepTRACE can map any feature-class relationship, regardless of whether the target behaviour is spatial, mobility-related, photometric, or a combination of these properties. DeepTRACE also supports import of arbitrary features outside its existing feature set (e.g. *FRET efficiency* for smFRET tracking which encodes relationships between particle motion and molecular conformation) if provided as a separate column in the tracking file; ensuring adaptability to future applications.

DeepTRACE requires cell boundary information to construct many of its powerful spatial features, and is currently optimised for rod-shaped bacterial cells. Future developments include support for other cell geometries; reinforcement learning via the *Track Inspector* tool, enabling the user to identify classification errors from which the model can learn to further boost performance on small datasets; and improved data mining capabilities of the *Discovery* tool.

In conclusion, DeepTRACE replaces rigid track analysis with a flexible, context-aware tool that learns and interprets the complex sequences of events captured by long single-molecule tracks. This approach enables the extraction of biological insights from minimal data, without requiring specialised ML hardware or expertise. Finally, DeepTRACE’s lack of hardcoded rules or pre-training, combined with its native handling of arbitrary input features and classes, makes it suitable for a wide range of applications and likely to remain relevant as experimental methods continue to evolve.

**Extended Data Figure 1.**
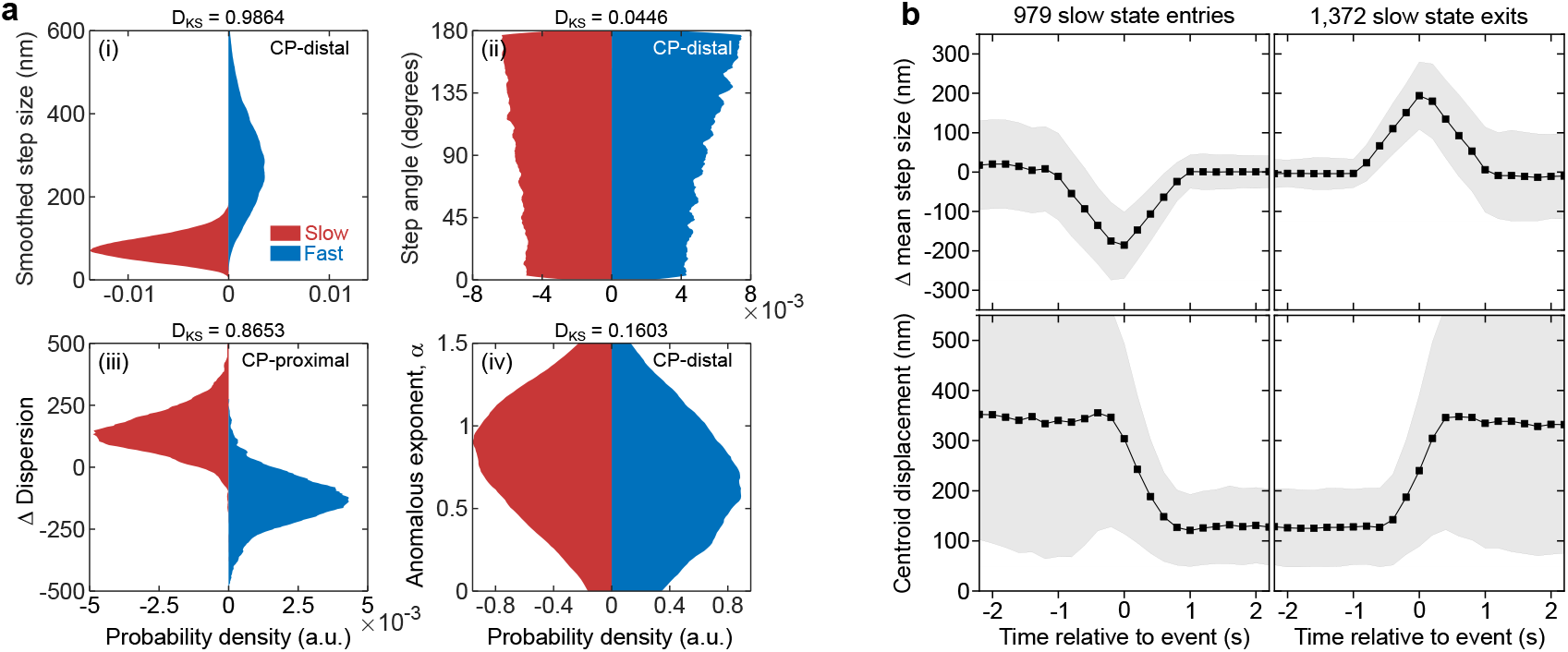
DeepTRACE visualisation of feature distributions. **a**, Class-conditional feature distribution plots for four features, showing the Kolmogorov-Smirnov statistic between classes (metric above plots) acting as a proxy for the static discriminatory power of each feature, produced using DeepTRACE’s *Feature Visualiser* tool. **b**, Mean values for two features (Δ *mean step size* and *centroid displacement*) relative to changepoints moving into (*N* = 979) and out of (*N* = 1,372) the slow diffusive state using DeepTRACE’s *Event Aligner* tool, revealing the temporal component of feature distributions. The shaded region indicates one standard deviation from the mean. Plots were produced using data from two-state reversible diffusion simulations (*D* = 0.01 *μ*m^2^*/*s and 0.2 *μ*m^2^*/*s) with ground truth for class assignment.

**Extended Data Figure 2.**
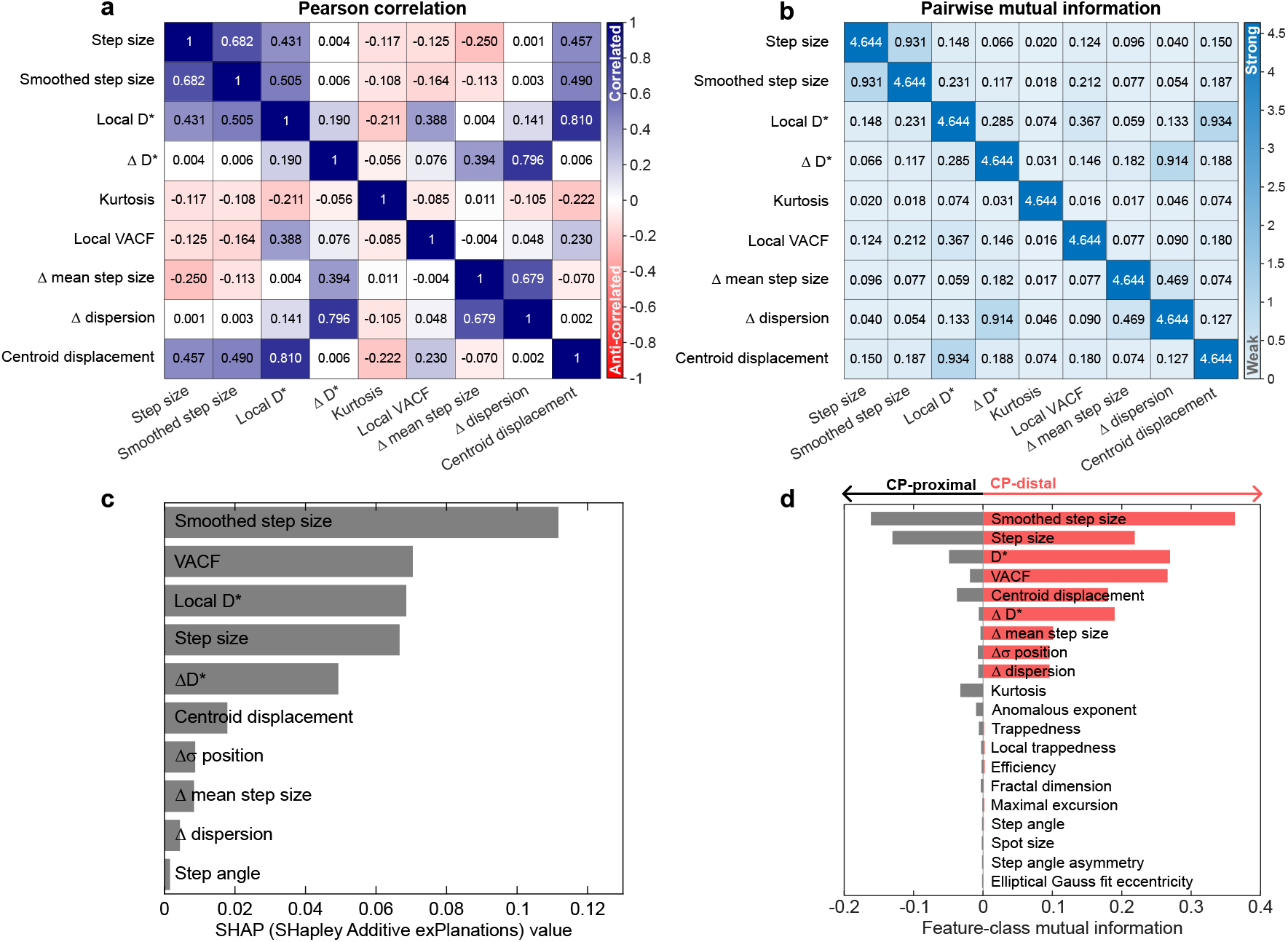
DeepTRACE feature relationships and ranking. **a**, Linear relationships between an example subset of features revealed by a Pearson correlation map between pair-wise combinations. **b**, Non-linear relationships between a subset of features, revealed by a mutual information map between pairwise feature combinations. **c**, SHAP (**SH**apley **A**dditive e**X**planation) values for features evaluated using a surrogate random forest model trained in DeepTRACE. **d**, Ranked mutual information between a subset of features and known class, providing a prediction of the relative static power of features, in both changepoint-proximal (defined here as the four localisations flanking each changepoint; grey bars, left) and changepoint-distal regions (red bars, right). All plots were produced using DeepTRACE’s *Feature Ranking* tool with data from two-state reversible diffusion simulations (*D* = 0.01 and 0.2 *μ*m^2^*/*s) using ground truth for class assignment.

**Extended Data Figure 3.**
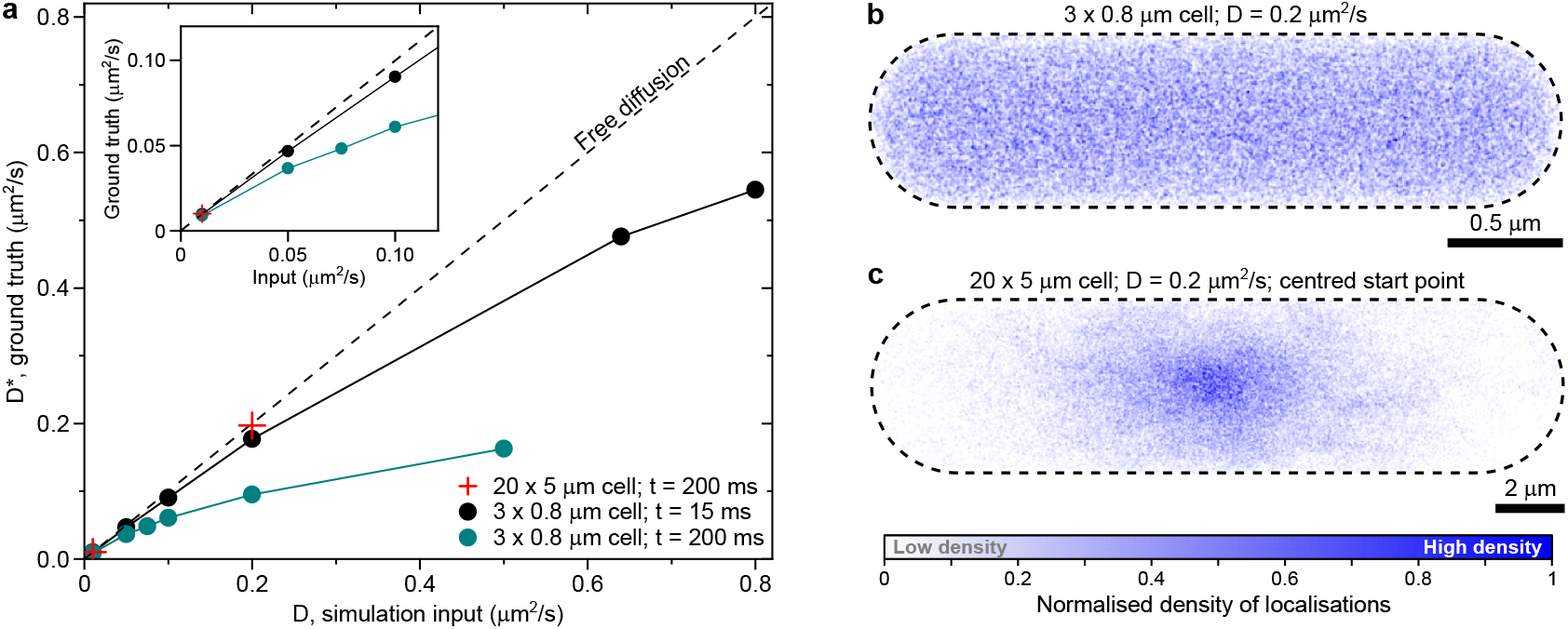
Anomalous sub-diffusion arises from cellular confinement. **a**, Apparent diffusion coefficients obtained from DeepTRACE’s *Diffusion Analysis* tool of simulated intracellular particle motion computed using MSD calculations of ground truth-segmented subtracks representing pure diffusive states; showing subdiffusive behaviour at both frame rates used (teal circles show timelapse imaging at intervals of 200 ms, black circles show continuous imaging at interframe times of 15 ms) resulting from confinement as molecules are influenced by the boundary. The dashed line represents free diffusion with an anomalous exponent of *α* = 1. Red crosses represent a control simulation showing greatly reduced anomalous sub-diffusion when particles are generated starting at the centre of an expanded 20 *μ*m × 5 *μ*m rod-shaped cell, illustrating the impact of confinement. **b-c**, Heatmap reconstructions of particle locations for all fast state track segments (*D* = 0.2 *μ*m^2^*/*s) using DeepTRACE’s *Spatial Mapping* tool with subtrack boundaries obtained from ground truth. **b**, All localisations from the fast diffusive state subtracks of 1,234 molecules simulated within a 3 *μ*m × 0.8 *μ*m rod-shaped cell, with uniformly distributed starting locations, showing a high degree of interaction with the enclosing cell membrane. The spatial map was constructed by positioning point spread functions (PSFs) with a full-width half maximum (FWHM) of 10 nm at each localisation and rendering into an image with a pixel scale of 5 nm. **c**, All localisations from the fast diffusive state subtracks of 490 tracks simulated within an expanded (20 *μ*m × 5 *μ*m) rod-shaped cell, with all molecules starting at the cell centre, showing a greatly reduced probability of interaction with the cell membrane. The spatial map was constructed using PSFs with FWHM of 50 nm rendered onto an image with a pixel scale of 25 nm.

**Extended Data Figure 4.**
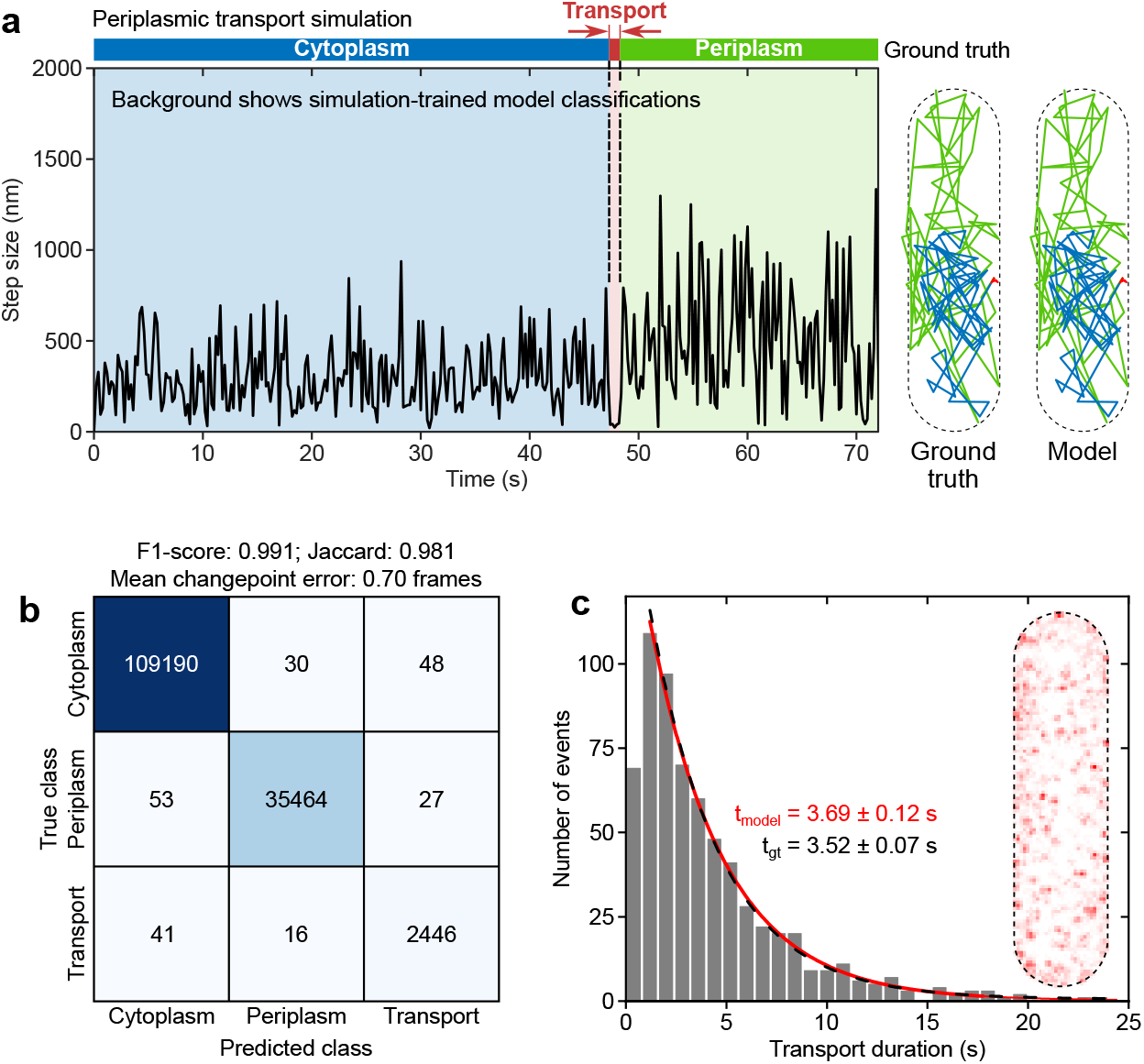
Simualtion of irreversible transport across the cytoplasmic-periplasmic membrane. **a**, Time series of the *step size* feature for a simulated membrane transport event using timelapse imaging, showing annotation from a simulation-trained model (inset colours), and the known ground truth (bar above). Shown to the right is the associated track plotted within the cell boundary (dashed line) coloured according to classifications of each timestep using DeepTRACE’s *Track Viewer* tool. **b**, Confusion matrix showing classification by a simulation-trained model of all localisations across all tracks in the test set, with performance quantified by macro-averaged F1-score, Jaccard, and mean changepoint error (statistics above). **c**, Residence times of all model-identified transport events using DeepTRACE’s *State Analyser* tool showing exponential fits to model identified residence times (red line), ground truth (dashed black line), and a heatmap of all timesteps classified by the model as undergoing transport (inset), using DeepTRACE’s *Spatial Mapping* tool. All tracks used for training were obtained from an entirely independent simulation to the evaluation dataset.

**Extended Data Figure 5.**
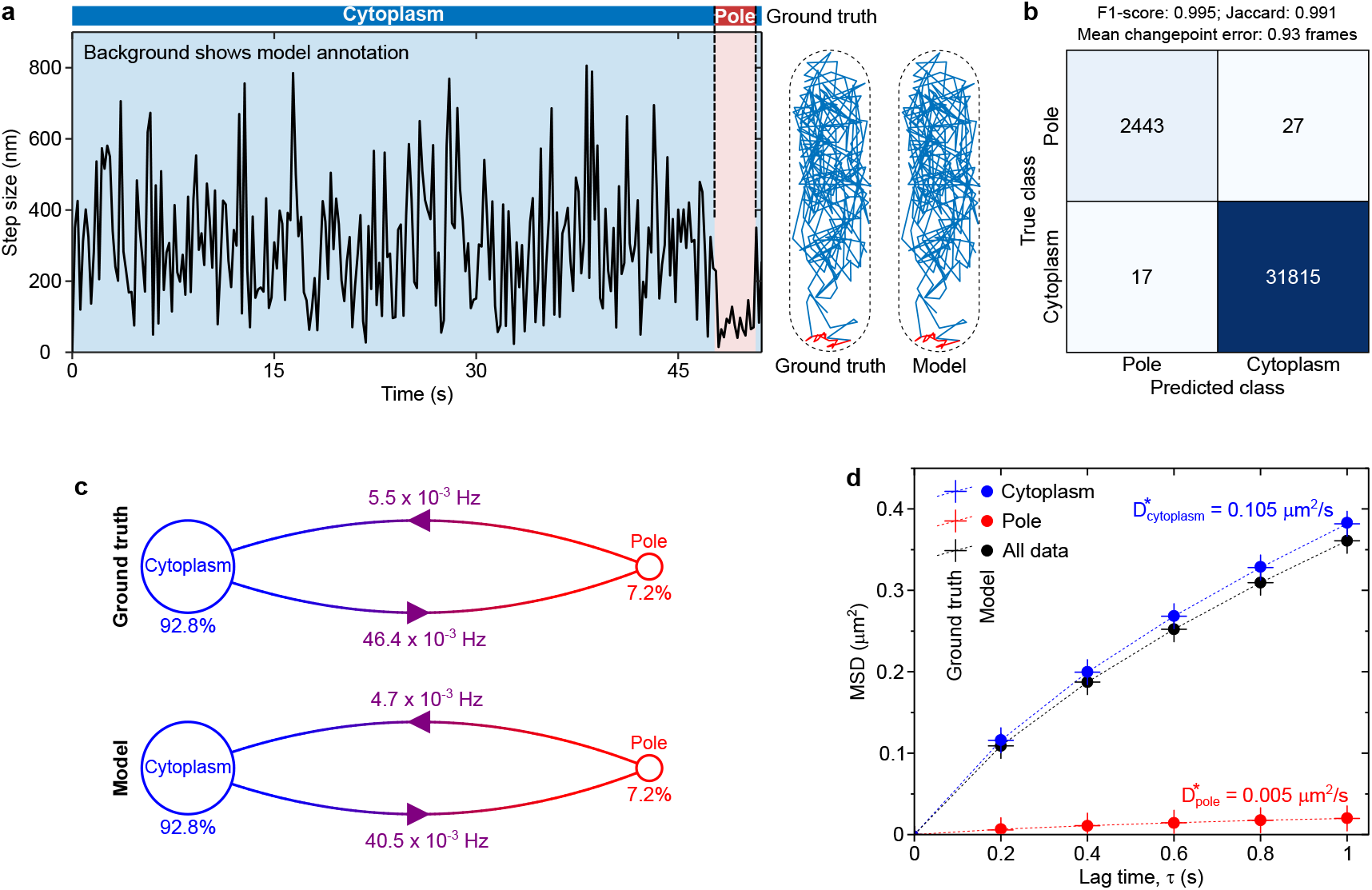
Simulation of reversible binding to a polar-localised plasmid. **a**, Example classification of a simulated molecular track displaying the evolution of the *step size* feature, with simulation-trained model annotation (inset colour), and ground truth (bar above) using DeepTRACE’s *Track Inspector* tool. Shown to the right is the associated track plotted within the cell boundary (dashed line), coloured according to ground truth and model classifications of each timestep using DeepTRACE’s *Track Inspector* tool. **b**, Confusion matrix showing classification by the simulation-trained model of all localisations across all tracks in the test set, with performance quantified by F1-score, Jaccard, and mean changepoint error (statistics above). **c**, Transition diagrams generated with DeepTRACE’s *Transition Viewer* tool for the model annotations and ground truth, showing occupancies of each class (circle areas and percentages), and transition rates (frequencies adjacent to arrows). **d**, MSD-lag time plots generated using DeepTRACE’s *Diffusion Analysis* tool for all model-segmented subtracks (blue and red circles indicate the cyptoplasmic and plasmid-bound polar states respectively), unsegmented tracks (black circles), and ground truth (crosses connected by dotted lines). The training dataset consisted of 1,249 tracks of which 1,061 were used for training and 188 for validation. Parameters were obtained from evaluation using a completely separate dataset consisting of 209 tracks, with a mean length of 164 localisations.

**Extended Data Figure 6.**
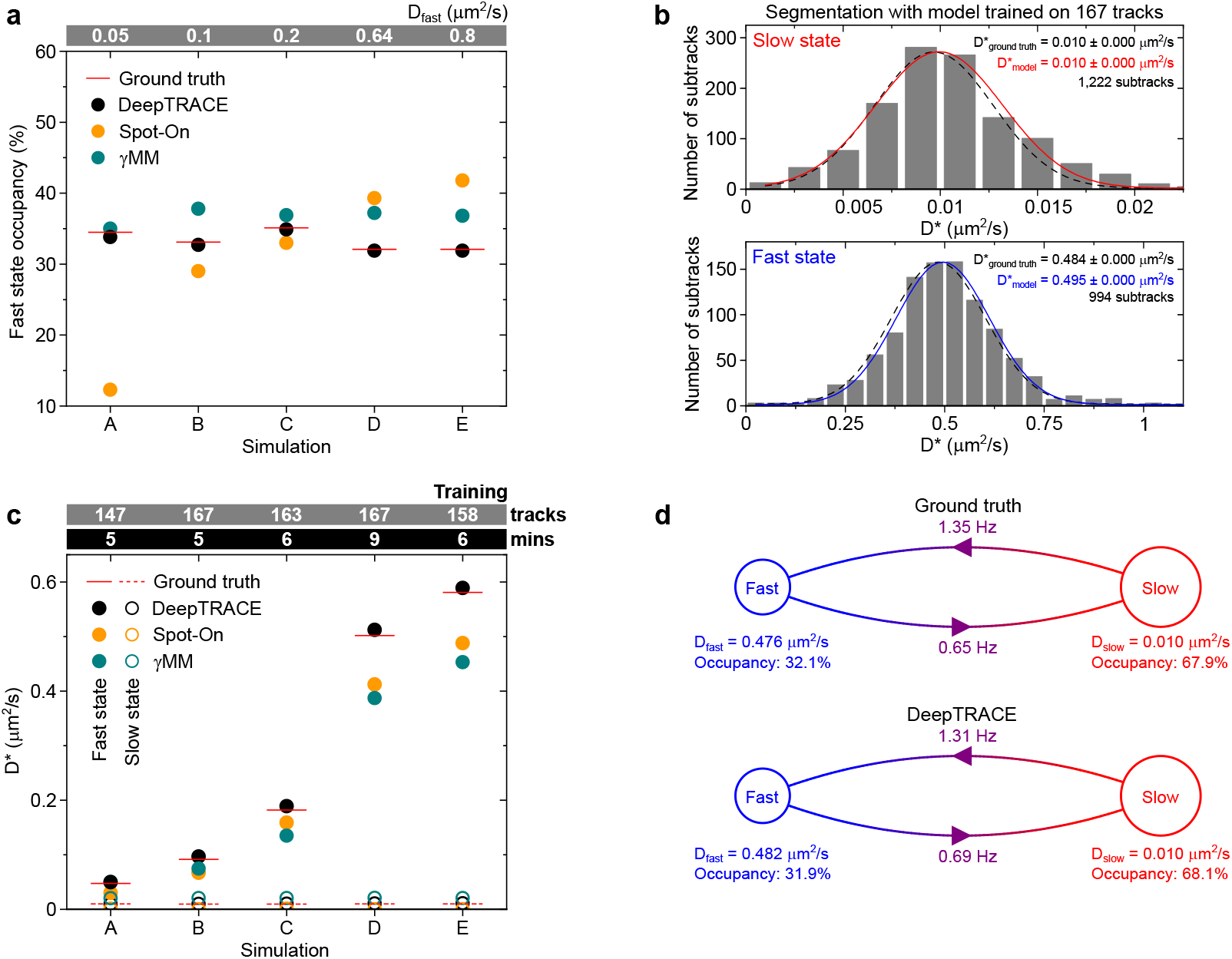
Performance comparison of DeepTRACE with established approaches. **a**, Comparison of DeepTRACE’s evaluation of state occupancies (black circles) to Spot-On (orange circles), γMM (teal circles), and ground truth obtained from MSD-lag time fitting of the same dataset using known segmentation boundaries (horizontal red lines). The bar above the plot identifies the input diffusion coefficient of the fast state, which is heavily reduced by cellular confinement, while the slow state is fixed at 0.01 *μ*m^2^*/*s. **b**, Histograms of apparent diffusion coefficient (*D*^∗^) for all model-segmented steps from a simulation of two-state reversible diffusion transitions with simulated input diffusion coefficients of 0.01 *μ*m^2^*/*s (slow state, top plot) and 0.64 *μ*m^2^*/*s (fast state, bottom plot), showing Gaussian fits to each population (red and blue lines). Shown inset are the fitted centre values, and overlaid on each plot is a dashed black line which represents the same fitting procedure performed using the ground truth segmentation boundaries (amplitude scaled for comparison). The classifying model was trained on 167 tracks. **c**, Performance comparison between DeepTRACE (black circles), Spot-On [21] (orange circles), and γMM (teal circles) for inference of diffusion coefficients across five simulations of reversible switching between diffusive states, compared to ground truth (red lines). DeepTRACE models were trained on fewer than 200 tracks and 10 minutes of processing on a laptop single CPU core (bars above). **d**, Transition diagrams generated with DeepTRACE’s *Transition Viewer* tool for the model annotations and ground truth, showing state occupancies (circle areas, and percentages), and transition rates (frequencies adjacent to arrows).

**Extended Data Figure 7.**
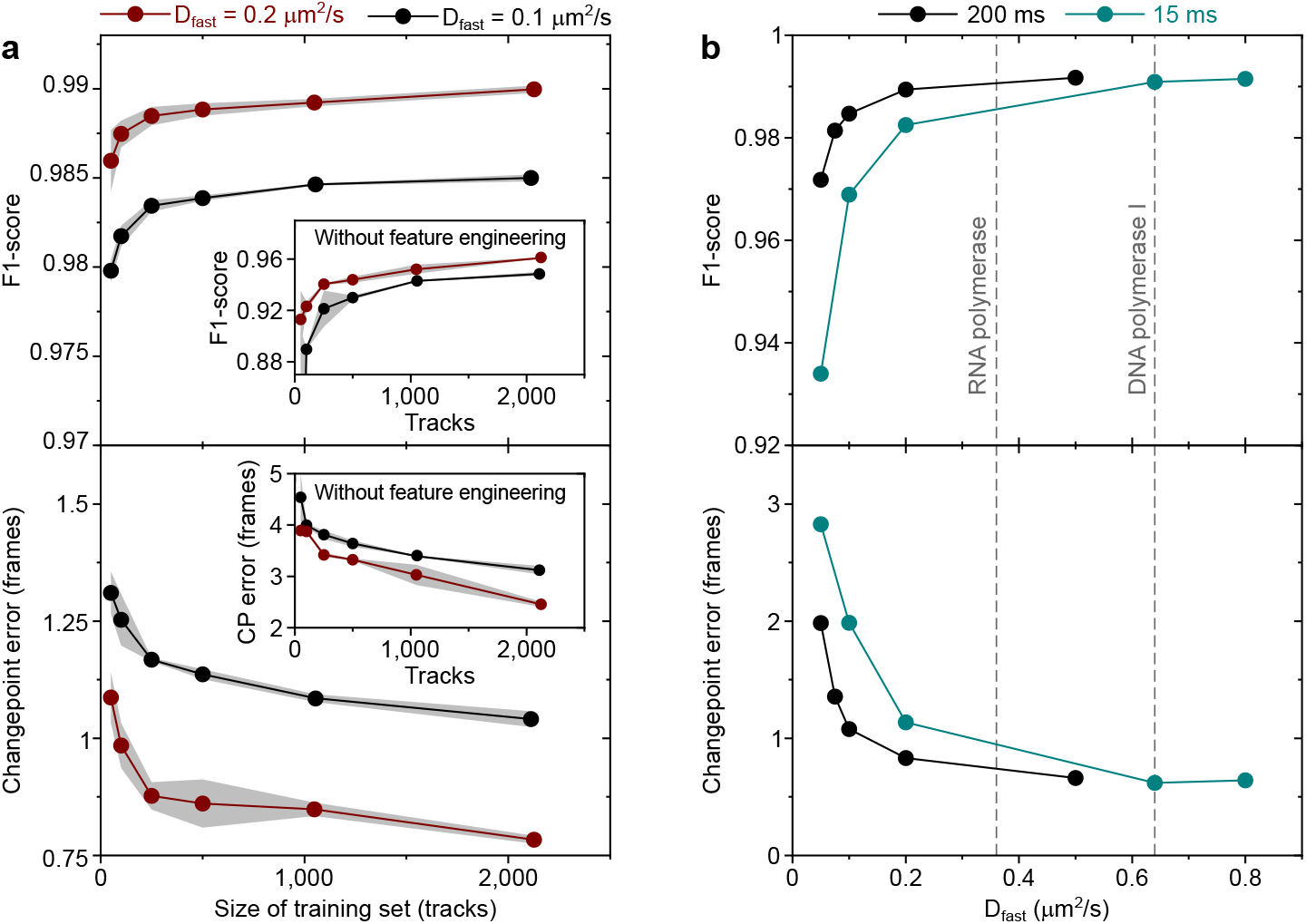
Classification performance of DeepTRACE models with varying training set size and separation of diffusion coefficients. **a**, F1-score (top) and mean changepoint error (bottom) as a function of training set size for simulations of timelapse imaging using intervals of 200 ms, with a slow state diffusion coefficient of 0.01 *μ*m^2^*/*s, and a fast state diffusion coefficient of 0.1 *μ*m^2^*/*s (burgundy circles) or 0.2 *μ*m^2^*/*s (black circles). The grey shaded region shows one standard deviation from the mean obtained from three independently-trained models. Shown inset are the same metrics obtained using identical parameters, model architecture, hardware, training data, and evaluation data, but eliminating the feature engineering process by restriction to only raw tracking data (*x, y, time*), performed twice for each datapoint. For all repeats, the training and validation split, shuffling, random track subsampling, and batch construction processes were performed independently using different random seeds. **b**, Variation of F1-score (top) and mean changepoint error (bottom) as a function of the diffusion coefficient of the simulated fast state, with the slow state fixed at 0.01 *μ*m^2^*/*s. The panels show performance from simulated timelapse imaging with an interframe time of 200 ms (black circles), and continuous imaging at 15 ms (teal circles) common with experimental techniques such as trackingPALM microscopy. Model training was restricted to a mean training set size of 1,084 tracks. The dashed reference lines indicate established diffusion coefficients of the target searching state for two DNA-binding proteins known to exhibit substantial slowdown due to intra-frame transient interactions with the chromosome (bacterial RNA polymerase [15] and DNA polymerase I [19]). For partial datasets (*N*_tracks_ ≤ 500), the original data were downsampled through random track elimination independently prior to splitting the training and validation data. Models were trained using a single BiLSTM layer with 100 hidden units, and a single attention head. All models were evaluated on data obtained from datasets generated independently from the training data.

**Extended Data Figure 8.**
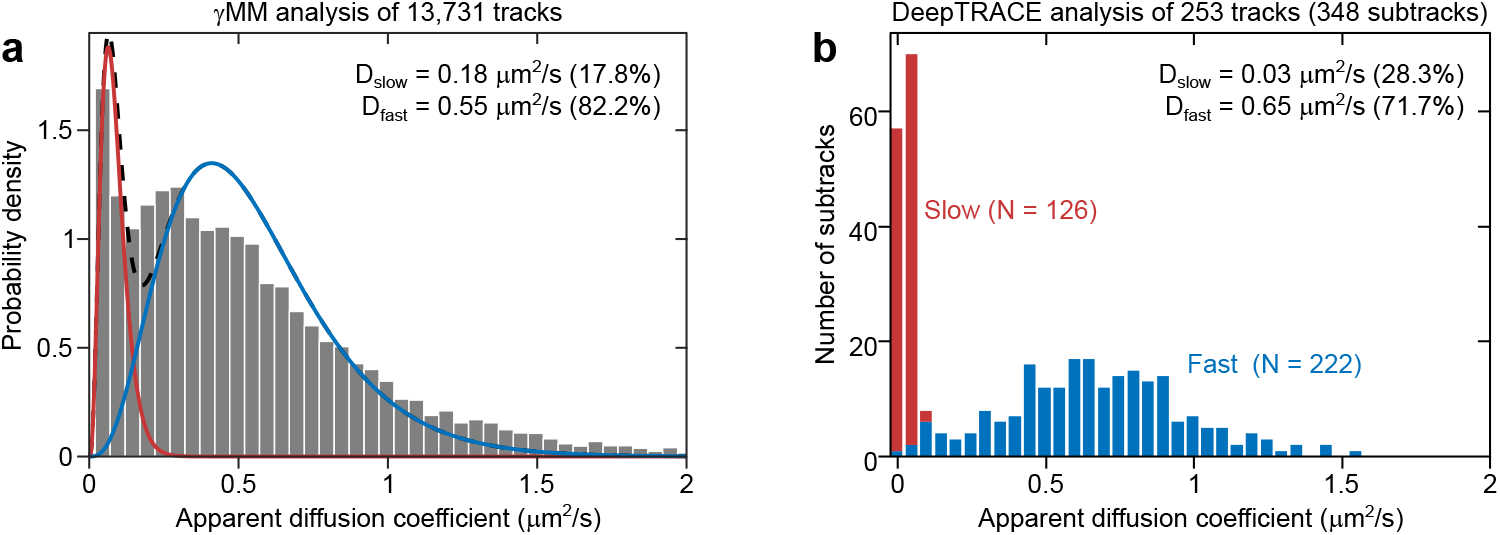
Improvement of diffusive state separation through track segmentation. Comparison using real single-molecule tracking experimental data of DNA repair by DNA polymerase I (Pol1) in *E. coli* recorded using 15 ms continuous imaging, five minutes after H_2_O_2_-induced DNA damage [24]. **a**, Diffusion histogram of tracking data compiled from MSD-lag time calculations of 13,731 Pol1 tracks truncated to the first 5 localisations, and analysed using a typical two-component Gamma mixture model (γMM) analysis [15, 19], showing the two fitted Gamma functions (red and blue solid lines) and their sum (dashed black line). **b**, A subset of 253 tracks selected from the same single field of view dataset by DeepTRACE based on track length and quality control criteria, segmented by a model into 348 subtracks, and compiled into a stacked histogram automatically using DeepTRACE’s *Diffusion Analysis* tool, illustrating improved state discrimination. The model was trained on 869 human-annotated tracks from separate experiments under similar experimental conditions.

**Extended Data Figure 9.**
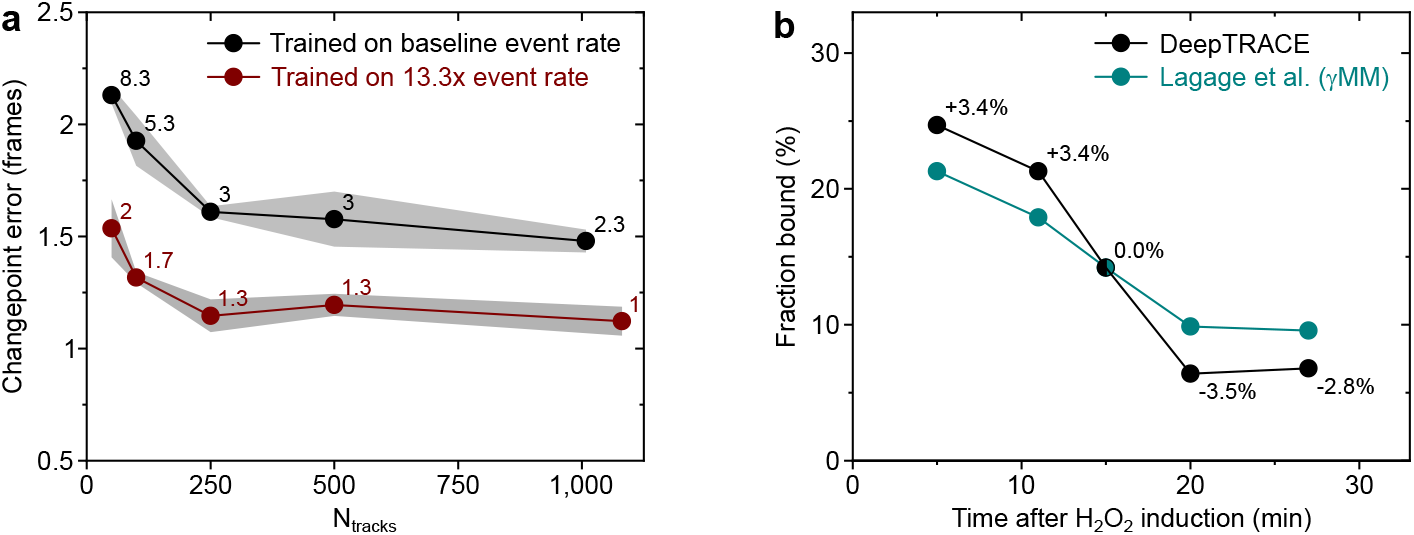
Training models using experimental perturbations in simulated and real experiments. **a**, Performance comparison of changepoint detection in reversible two-state diffusion simulations, between models trained on data with class transition rates matching the evaluated dataset (0.1 Hz and 0.05 Hz, black circles), and models trained with data containing a perturbation enhancing rates to 1.33 Hz and 0.67 Hz (red circles). The changepoint error is the mean number of frames separating the predicted and known changepoints from the simulation ground truth (see **Online Methods**). The evaluated dataset contained 118 tracks with 43 change-points. Points and shaded areas show the mean and standard deviation of the changepoint error respectively for three separately trained models, each using an independently randomised training/validation split and shuffled batches. Data point labels show the mean number of unmatched changepoints. Both training datasets were generated independently of the data used for evaluation; with each downsampled subset (N_tracks_ ≤ 500) obtained by a separate resampling of the full simulation by random track elimination (from 1,186 and 1,272 tracks in the original baseline and enhanced simulations respectively). **b**, Fraction of time spent by DNA polymerase I molecules in the DNA-bound state during a timecourse series of real single-molecule tracking experiments following exposure to the DNA-damaging agent H_2_O_2_, as evaluated by a model trained on 453 human-annotated tracks from a separate experiment 4 min following H_2_O_2_ exposure (black circles), compared to estimates obtained from γMM analysis of the same recordings as reported by Lagage et al. [24] (teal circles). While no ground truth is available for real experimental data, both analyses reflect the expected reduction in bound state occupancy over time. Data labels indicate the difference in bound population between estimates from DeepTRACE and γMM.

**Extended Data Figure 10.**
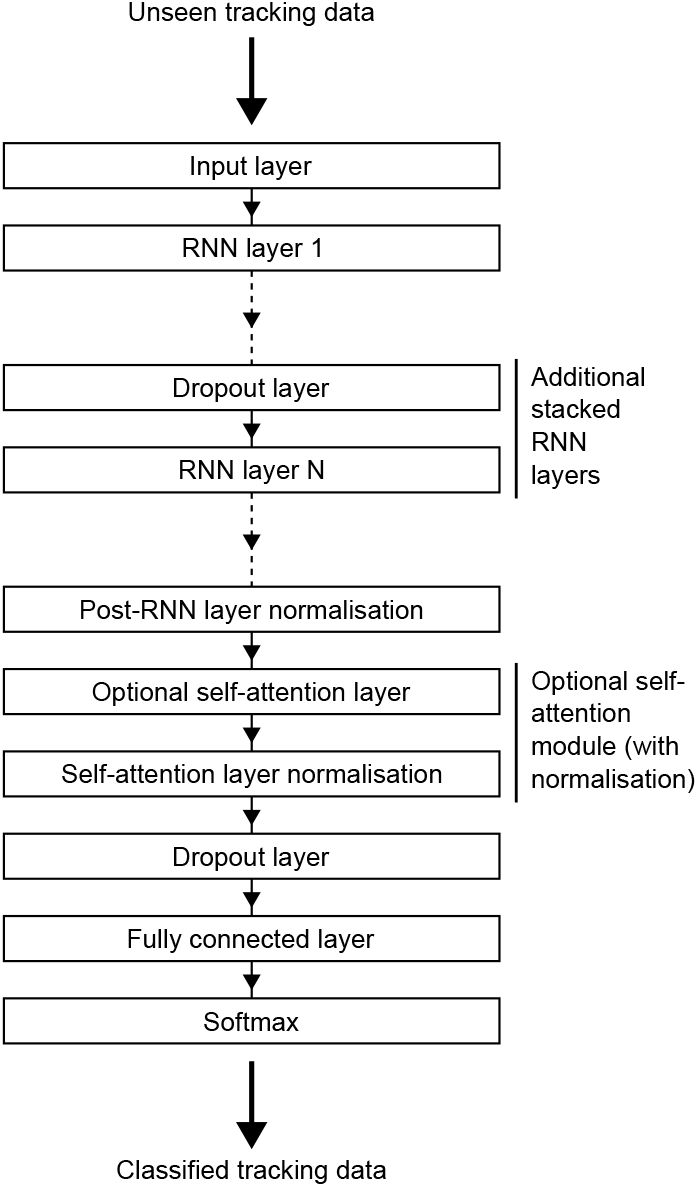
Modular architecture for RNN-based models. RNN-based DeepTRACE models can be constructed by the user through the GUI using a standardised model architecture based on a deep sequence model design incorporating an arbitrary number of stacked RNN layers, and optional self-attention. The model architecture performs sequence-to-sequence classification of unseen data.

**Extended Data Table 1.** Performance comparison of estimated kinetic parameters from 1,292 simulated tracks using a DeepTRACE model trained on ground truth labels from 167 tracks in an independent simulation (**Sim-GT**), a DeepTRACE model trained on human annotations of the same 167 tracks (**Sim-Human**), and a DeepTRACE model trained on human annotations of 453 tracks from real experiments of DNA polymerase I (Pol1) repairing H_2_O_2_-induced DNA damage (**Real-Human**), with the same 1,292 simulated tracks processed using the popular software **Spot-On**, and analysis using a Gamma mixture model (γ**MM**). Tracks were generated using a reversible two-state diffusion simulations for continuous imaging with 15 ms exposures, and diffusion coefficients of 0.01 *μ*m^2^*/*s and 0.64 *μ*m^2^*/*s, matching reported Pol1 mobilities in *E*.*coli* [24]. The discrepancy between the ground truth apparent diffusion coefficient and the simulation input values — which is most prominent in the fast state — arises primarily from cellular confinement. All training data were generated independently from the dataset used for evaluation.

| Parameter | Simulation input | Ground truth | DeepTRACE Sim-GT | DeepTRACE Sim-Human | DeepTRACE Real-Human | Spot-On | γMM |
| --- | --- | --- | --- | --- | --- | --- | --- |
| Slow state occupancy (%) | 66.7 | 67.9 | **68.1** | 68.5 | **68.1** | 60.7 | 62.8 |
| $D_{\text{slow}}^*$ ($\mu\text{m}^2/\text{s}$) | 0.0100 | 0.0098 | **0.0105** | **0.0105** | 0.0107 | 0.000 | 0.021 |
| $D_{\text{fast}}^*$ ($\mu\text{m}^2/\text{s}$) | 0.0640 | 0.5017 | 0.5117 | **0.5050** | 0.5145 | 0.412 | 0.387 |
| Slow → Fast transition rate (Hz) | 0.667 | 0.654 | **0.649** | **0.649** | 0.684 | N/A | N/A |
| Fast → Slow transition rate (Hz) | 1.333 | 1.352 | 1.304 | **1.379** | 1.243 | N/A | N/A |

## Online Methods

### DNA polymerase I experimental data

Previously published single-molecule fluorescence tracking experimental data of DNA polymerase was kindly provided by Lagage et al [24] in the form of TIF-formatted fluorescence video files of single-molecule tracking under continuous imaging with an exposure time of 15 ms, brightfield reference images, and MicrobeTracker-formatted cell segmentations. Fluorescence excitation of the organic dye JF549, covalently linked to DNA polymerase I via a HaloTag linker, was performed using variable angle epifluorescence excitation with a 561 nm laser. Under-labelling ensured temporally separated single-molecule tracks. In each case we used the first data acquisition following H_2_O_2_ exposure (4 − 7 mins) for model training.

### Simulations

Simulations of single-molecule tracking experiments were generated using the Brownian dynamics simulator Smoldyn, with synthetic microscopy video sequences rendered using the software SMeagol.

Smoldyn simulations were performed inside an idealised rod-shaped cell, modelled as a cylinder (length 2.2 *μ*m, radius 0.4 *μ*m), and enclosed at both ends with hemispherical end caps of radius 0.4 *μ*m (total length 3 *μ*m). The surrounding simulation volume consisted of a regular cube of side 5 *μ*m. 3D coordinates and diffusive states were computed at 1 ms and 10 ms intervals for inter-frame times of 15 ms and 200 ms respectively.

In both real and simulated data, changepoints were not synchronised to camera shutter actuations; consequently the vast majority of localisations immediately preceeding changepoints comprise a mixture of states. This lack of synchronicity in both real and simulated data unavoidably contributes to the changepoint error, and we also note the existence of a class-asymmetry artefact that arises from the classification of different diffusion coefficients with sub-frame changepoints which further complicates classification. To assign a single ground truth class from the constituent subframes containing mixed classes we therefore explored several approaches, including the use of median, modal, and maximum diffusion values; settling on a ground truth definition as the state of the first subframe of each frame.

Fluorescence frames were generated by SMeagol from the coordinates and diffusive states exported by Smoldyn, using integration of fluorescence emission over 10 ms, at intervals of 15 ms to match common continuous imaging frame rates, and intervals of 200 ms for timelapse imaging. Simulated photoactivation ensured that only one molecule is active at any given time, and fluorescence emission continued until permanent photobleaching. Fluorescence intensities were sampled from a Gaussian distribution, with a simulated 2D Gaussian point-spread function (*σ* = 150 nm). Poisson-distributed background noise and camera simulation parameters (e.g. gain) were tuned to reproduce imaging data comparable to a typical super-resolution fluorescence microscope. Photo-bleaching times for each experiment were uniformly distributed across the range 65 − 150 frames. The photobleaching time for each molecule in the simulation was then drawn from a Gaussian distribution centred around this time.

Videos for each molecule were constructed in stacks of 500 frames. These videos were stitched together spaced by ten empty frames for continuous imaging and five empty frames for timelapse imaging, producing aggregated video sequences of 25,245 and 25,490 frames respectively. Datasets for all simulations were generated in independent batches of 300 and 2,000 molecules. To explore the variation of model performance with dataset size, training datasets were reduced by downsampling via random track elimination, and expanded by concatenation.

Three biological scenarios were simulated: (i) reversible interconversion between two diffusive states, (ii) binding to a polar-localised plasmid, and (iii) irreversible transport across the cytoplasmic-periplasmic membrane.

#### (i) Reversible interconversion between two diffusive states

Two-state simulations consisted of a slow state with a fixed diffusion coefficient of 0.01 *μ*m^2^*/s*, and a fast state that was varied between simulations across the range 0.05 *μ*m^2^*/s* to 0.8 *μ*m^2^*/s*. Each molecule began at a random position within the cell cytoplasm, undergoing Brownian motion, and confined by the cell membrane with reflective boundaries. State occupancies began in equilibrium, with two-thirds of molecules beginning in the slow state. For timelapse imaging, transition rates between diffusive states were set at 0.1 Hz and 0.05 Hz for fast-to-slow and slow-to-fast transitions respectively; an asymmetry that simulated class imbalance. For simulations of continuous imaging, the transition rates were adjusted to 0.667 Hz and 1.333 Hz to produce a similar density of observed changepoints in the faster photobleaching regime. This scaling of transition rates reflects the increasingly common use of temporal patterning techniques and improved probe stability that enables the experimentalist to adjust the imaging timescale to capture reasonable densities of transitions (e.g. extending dark times for slower processes). These class transition rates were varied only for simulations investigating the impact of experimental perturbation on model performance (**Extended Data Fig. 9a**); in which class transition rates under continuous imaging were reduced to 0.1 Hz and 0.05 Hz under continuous imaging for the evaluation dataset.

#### (ii) Reversible binding to a slow diffusing polar-localised plasmid

A molecule was simulated with a diffusion coefficient of 0.2 *μ*m^2^*/s* diffusing freely within the cell cytoplasm, and able to bind reversibly to a polar-localised plasmid within each end cap (a region defined as 200 nm from the cell pole). Within these regions the molecule experienced a binding probability equivalent to an on-rate of 0.1 Hz. Once in the bound state, the molecule was confined to the polar region, with diffusive state 0.01 *μ*m^2^*/s*, until released from the plasmid with an off rate of 0.05 Hz, and transitioned back to the initial diffusive state in which it was again able explore the entire cytoplasm. This process was allowed to repeat multiple times before probe photobleaching. Simulations were initiated in equilibrium, with 4.07% of molecules initially in the polar-bound state with the remainder beginning in the cytoplasm; a fraction that accounted for both transition rates and the relative volume fraction within which transitions to the polar state can occur. In each case molecule start locations were drawn from a uniform distribution within the permitted volume regions for each class.

#### (iii) Irreversible transport across the periplasmic membrane

Transport across the membrane separating the cytoplasm and periplasm (a 20 nm-thick shell at the cell periphery enclosing the cytoplasm with reflective boundaries) was simulated for a single molecule initially undergoing free diffusion within the cytoplasm with a diffusion coefficient of 0.2 *μ*m^2^*/s*. Collision with the membrane resulted in a probability of transmembrane transport set by an ‘absorption rate’ of 0.001 Hz, causing it to become immobilised at the closest point of contact. The molecule was otherwise reflected at the boundary. Following transport, the molecule was transferred irreversibly to the periplasmic space, which it continued to explore with a diffusion coefficient of 1.0 *μ*m^2^*/s*.

### Single molecule localisation and tracking

Candidate single molecule locations were obtained from spatial bandpass filtering of each fluorescence video frame, using a threshold tuned to the imaging conditions. Super-resolution localisation was then performed through 2D elliptical Gaussian fitting of candidate positions, producing several features (e.g. *peak intensity, width of Gaussian fit major axis*) that were incorporated into DeepTRACE’s raw feature set. Localisations were connected into tracks using a nearest-neighbour search between neighbouring frames within a maximum distance of 7 pixels (672 nm). Tracking was performed independently within each segmented cell boundary. For simulated data the maximum connecting distance for tracking was extended to 50 pixels (4.8 *μ*m) as false track connections from temporal overlap were not possible due to the sequential nature of the simulations. A tracking memory parameter of one was used, allowing track connection to bridge single frame blinking events.

### Track filtering

Tracks imported into DeepTRACE were filtered with a minimum track length of 50 and 65 frames for simulated and experimental data respectively. After filtering for overall length, tracks were processed using DeepTRACE’s ‘*truncation by localisation filtering* ‘ option with a tolerance of two localisations, resulting in any tracks that coincide with more than two localisations (regardless of whether they were tracked) being truncated to eliminate any temporal overlaps for which track history is uncertain. This information-preserving approach can result in tracks shorter than the minimum track length appearing in the dataset.

### DeepTRACE graphical interface

All analysis was performed using DeepTRACE’s cross-platform Graphical User Interface (GUI). The GUI supports data pre-processing, feature engineering and selection, human annotation, model construction and training, track segmentation, and a wide range of downstream analytics tools.

### Feature engineering

Data preprocessing by DeepTRACE engineered 60 features. These features have varying uses in different experimental contexts. We group these features broadly into four main categories,

- **Raw features**: Features taken directly from the source data (localisation fitting parameters, photometric properties from images, and tracking data), such as *peak intensity, x-y coordinates*, or *local background noise*.
- **Static engineered features**: Features that are engineered without temporal context, and depend only upon the current localisation or two points immediately flanking the current localisation. Examples include *step size, distance to the nearest cell membrane*, and *step angle*.
- **Local window features**: Features that are computed from a local context window, such as *smoothed step size, step angle asymmetry*, or *path straightness*.
- **Delta features**: Features that are computed from comparison of two local context windows, immediately preceding and following the datapoint. These features are particularly powerful for the identification of changepoints, especially when implemented for small models and those without native temporal context. In larger models, these simplified representations of temporal behaviour accelerate learning by shortcutting the process of constructing higher order representations of changepoints from raw features. Delta features are computed as either the difference or the ratio between the window pair, with the preceding window defined as the denominator or subtrahend. Examples include Δ*dispersion, mean step size ratio*, and *centroid displacement*.

As window size is a trade-off between accuracy and sensitivity to changepoints, for all local features except for *smoothed step size* we used a local window size of 11 frames (five localisation flanking the classified localisation); this scale was obtained from empirical testing with a binary search. All delta features used a pair of windows each containing four frames, with the evaluated localisation being the first datapoint of the second temporal window.

Close to the track end points in which one frame becomes compressed, the value for each feature is scaled appropriately where possible by the available number of accessible frames, trading increased noise in exchange for improved temporal sensitivity and access to end point proximal regions. For some static features it is not possible to scale values when windows are compressed close to end points (e.g. *relative step angle*, which requires a minimum of three localisations). As data imputation in many experimental tasks can produce errors arising from temporal context, DeepTRACE instead sets these values to zero, and we used options accessible within the GUI to truncate the first and final two localisations prior to training, as described in the section ‘Preparation of training data’.

We provide here formal definitions for non-intuitive features, and conceptually simple features which may have ambiguous definitions such as *smoothed step size*. Remaining features for which the definition is obvious (such as *step size* which computes the simple Euclidean distance between frames, or *time step* which computes the temporal separation between localisations to learn behaviours associated with single-molecule blinking events) are listed separately.

### Centroid displacement

*Centroid displacement* is a measure of the Euclidean distance between centroids of localisations in windows either side of the localisation being evaluated. Specifically, *centroid displacement* (*d*_*C*_) was defined as,

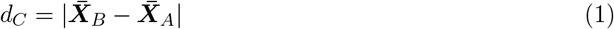

Where 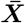 represents the centroid of all *W* localisations within the preceding (*A*) and following *(B)* windows, defined as follows,

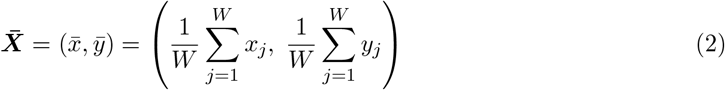

### Delta dispersion

*Delta dispersion* quantifies changes in the spatial spread of localisations between two adjacent temporal windows. We defined the dispersion *ψ* of a window containing *W* positions *X*_*i*_ = (*x*_*j*_, *y*_*j*_) as the mean Euclidean norm per point, summed over all points, relative to the window centroid,

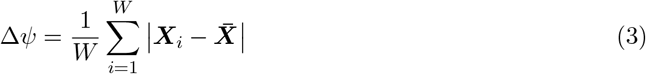

*Delta dispersion* and *dispersion ratio* were then computed by the difference or ratio of *ψ* within each window.

### Smoothed step size

*Smoothed step size* is particularly useful for suppressing localisation error, statistical noise, and projection effects. DeepTRACE provides multiple smoothing methods, including Gaussian kernel smoothing, moving mean, local regression such as (R)LOWESS and (R)LOESS, and the Savitzky-Golay filter. For all work presented here, a median filter with a window size of 3 was used.

### Longitude and latitude

To map localisations onto a universal model-cell coordinate system, we perform a transformation along the major and minor axes, which we term *longitude* (*L*), and *latitude* (*ℓ*). The *longitude* feature is obtained from the contour length of the projected position along the cell midline, while *latitude* is computed as the relative distance between the projected midline location and cell boundary. The coordinate system is designed such that the origin is placed at the mid-point of the midline contour, with the boundary at ±0.5. The midline vertices, *M*_*i*_, are obtained by averaging each left and right coordinate pair from the MicrobeTracker-formatted boundary,

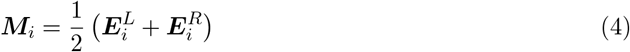

Where 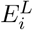 and 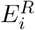 represent the paired vertices defining each edge of the segmented cell boundary. Each localisation is projected onto the closest midline segment [*M*_*i*_, *M*_*i*+1_], and *longitude* is computed as,

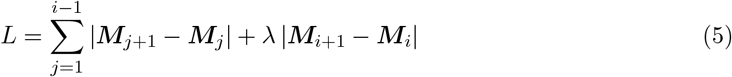

Where *λ* is the fractional position along the segment,

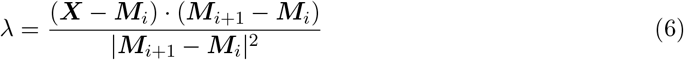

Finally, we define *latitude* as the relative perpendicular distance from the midline to the cell boundary,

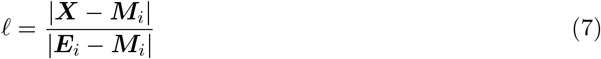

Note that both *latitude* and *longitude* can exceed the cell boundary if the input tracking data associates such localisations with the segmented cell; this enables DeepTRACE to retain tracking of localisations that fall outside of the segmented boundary due to localisation uncertainty or drift, and to process nearby contextual data, while also being able to exclude such data when not desired, for example in constructing spatial maps.

### Local straightness

Path *straightness* is a geometric local feature quantifying the directness of the molecule’s motion. *Straightness* was defined as the ratio of the distance between start and end points of the windowed track to its full contour length,

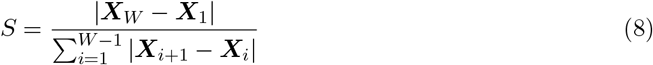

### Local path efficiency

*Efficiency* is a normalised version of path *straightness*. We used the definition of *efficiency* put forward by Kowalek et al. [25]; for a local window containing *W* localisations, path *efficiency* was computed as the ratio of the squared total displacement (between window start and end points) to the product of the number of steps and the sum of the squared step lengths,

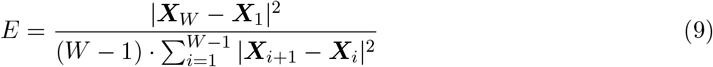

*E* → 1 for directed motion, and *E* → 0 for extremely convoluted paths. The normalisation of path *efficiency* makes it more resistant to compressed data sampling regions, such as towards the ends of tracks. While dynamically compressing the local window increases noise, it avoids reporting on an artificial increase in the directness of the path towards the ends of the track and in regions of excessive photoblinking.

### Local fractal dimension

We used the definition of *fractal dimension* (*D*_*f*_) provided by Katz and George [26] as follows,

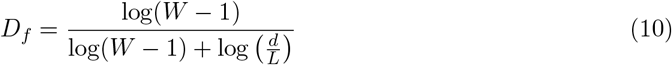

Where *W* is the total number of localisations in the window, *d* is the maximum distance between any two points in the window, and *L* is the total contour length. The feature has been shown to be effective at discriminating mobility modes.

### Local trappedness

Following Saxton et al. [27] the *trappedness* was defined as,

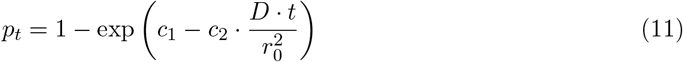

Where *D* is the diffusion coefficient obtained from a linear fit to the mean squared displacement with a maximum lag time, *t*, of two frames, *r*_0_ defines the trapping region represented by half of the maximum step size along the path, and *c*_1_ and *c*_2_ are empirical fitting constants adjusted accordingly for our experiments.

### Local step angle asymmetry

*Step angle asymmetry* (*A*_*s*_) is a proxy for the degree of spatial confinement. Expanding on the definition from Izeddin et al. [28], we defined the *step size asymmetry* as,

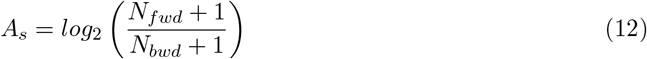

Where *N*_fwd_ represents the number of forward steps with step angles between 0 and 30 degrees, and *N*_bwd_ represents the number of backward steps with step angles between 150 and 180 degrees. Values for *A*_*s*_ are negative for backward step angle biases, and positive for forward biases. We retained the log_2_ notation for consistency with past work [28] despite the rescaling of this feature to a symmetric distribution around zero being unnecessary due to the Z-score feature scaling internal to DeepTRACE. The introduction of +1 here improves robustness of the metric by preventing division or multiplication by zero when operating on small local windows.

### Local kurtosis

The gyration tensor for a set of points *X*, within a window containing *W* localisations was defined as,

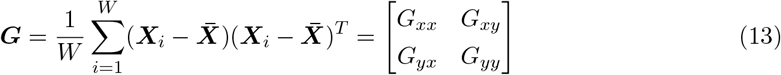

The dominant eigenvector, representing the largest spread of localisations, was obtained by diagonalirsing ***Gv*** = *λ****v***, where *λ* are the eigenvalues, and *ν* the eigenvectors defining the principal axis. The projection, *p*_*i*_, of each point along the dominant eigenvector *ν*_*max*_ was obtained by,

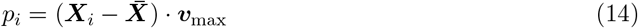

The local *kurtosis* is then computed from these projected positions as,

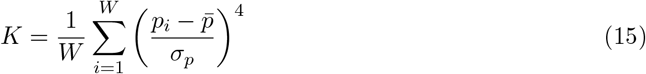

Where *σ*_*p*_ represents the standard deviation of positions projected along the dominant eigen-vector.

### Local step size kurtosis

*Step size kurtosis* is a measure of tailed-ness of the step size distribution, defined as,

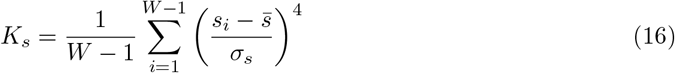

Where *s*_*i*_ = ||***X***_*i*+1_–***X***_*i*_|| is the size of the *i*^th^ step, 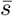 is the mean step size, and *σ*_*s*_ is the standard deviation of step sizes between the *W* localisations within the local window,

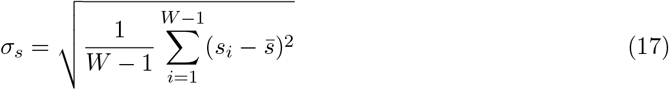

### Local maximal excursion

Local *maximal excursion* (*ME*) quantifies the extent of large, isolated displacements within a local window relative to the net displacement over that window. It is particularly effective at detecting transient high-mobility events, such as sudden or transient escape from confinement. The local *maximal excursion* for a window centred at localisation *i*, was computed as the ratio of the maximum Euclidean distance between consecutive points within the window to the net displacement across the window,

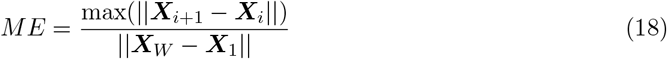

Where ***X***_1_ and ***X***_*W*_ are the spatial coordinates of the molecule in the first and last frame of the local window respectively. When combined with features that encode local diffusion coefficient estimates, the model can be particularly effective in identifying rapid or brief transitions in diffusive state.

### Local Velocity Autocorrelation Function (VACF)

The local *velocity autocorrelation function* (*V*) quantifies the temporal correlation of motion over short time scales, capturing the tendency of the molecule to maintain its direction of movement. It was computed as the average dot product between a step vector and a step vector offset by fixed temporal distance (*n* frames) relative to itself, as follows,

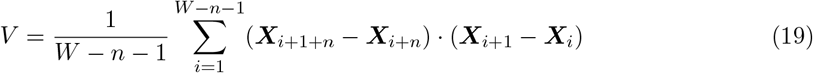

Throughout this work we used *n* = 1. Positive values indicate persistent motion (the molecule tends to continue in the same direction), while negative values indicate anti-persistent motion (the molecule tends to reverse direction). Values near zero correspond to random motion. The local *velocity autocorrelation function* is particularly useful for assisting models in identifying changes in diffusion type.

### Spot size and area

*Spot size* (*ϕ*) and *spot area* (*A*) quantify the spatial extent of localisations which encodes information about intra-frame particle motion, particularly in continuous exposure experiments. These features were computed from the standard deviation of the major and minor axes (*σ*) of the 2D elliptical Gaussian fits during the localisation process,

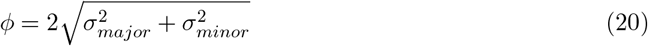

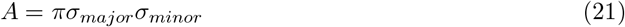

### Step angle features

Three *step angle* features were engineered: the relative step angle between successive steps (*θ*_*rel*_ step angle relative to the image (*θ*_*img*_), and step angle relative to the major cell axis (*θ*_*cell*_), using the step vectors at positions *i* − 2 to *i*,

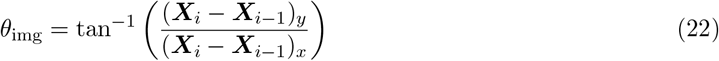

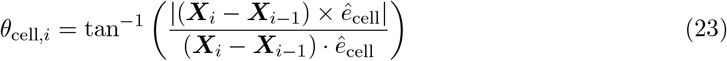

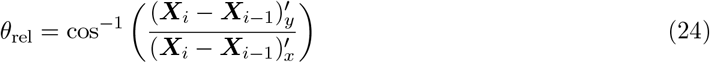

Where *ê*_*cell*_ is a unit vector along the major cell axis, and (***X***_*i*_ − ***X***_*i*−1_)^′^ represents the step vector from the current step rotated onto the frame of the previous step using the transformation 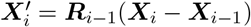; the rotation matrix *R*_*i*−1_ is defined as,

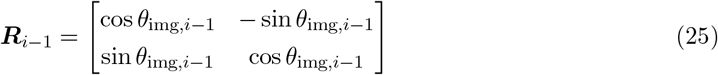

Note that for *relative step angle*, the previous angle (*θ*_*img*,*i*−1_) requires the localisation in frame (*i* − 2) to compute the previous step vector, and as a result the *relative step angle* feature contains two zero values at the start of each track due to insufficient data.

### Apparent diffusion coefficient

Using a mean squared displacement (MSD) matrix compiled for all pairwise displacements within the window, the local *apparent diffusion coefficient* (*D*^∗^) feature was computed from a linear fit to,

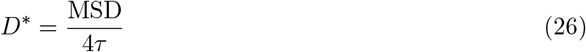

Where *τ* is the lag time. For all models trained here, we chose to use a maximum lag time equal to the window size to simplify the definition of local features, something not recommended in most contexts. This approach differs significantly from our formal calculation of *D*^∗^ used for global diffusion analysis in which calculation is performed using *τ*_*max*_ = 2.

### Feature selection

Feature selection was performed in five stages,

1. Initial feature selection based on experimental domain knowledge.
2. Visual screening of class-conditional feature distributions and class separability metrics using DeepTRACE’s *Feature Visualisation* tool to identify the static discriminatory power of each feature.
3. Feature ranking via feature-class mutual information using DeepTRACE’s *Feature Ranking* tool.
4. Redundancy reduction through pairwise Pearson correlation thresholding (|*r*| *<* 0.9), and identification of non-linear dependencies using mutual information heatmaps, both computed using DeepTRACE’s *Feature Ranking* tool.
5. Final selection used permutation importance measured on a candidate model in changepoint proximal and distal regions, via DeepTRACE’s *Permutation Importance* tool.

Redundant features were removed iteratively based on feature-class mutual information, permutation importance, and class separability metrics, in order of increasing utility until no feature redundancies remained.

In practice, because most of our experiments involved systematic comparisons of model performance under varying conditions (e.g., training set size, separation of diffusive states, experimental perturbation), it was critical to minimise variability introduced by the feature selection itself arising from a large input feature space; we therefore further constrained the feature set to a small, high-confidence subset to ensure stable and reproducible learning dynamics across runs, resulting in fairer comparisons.

DeepTRACE’s SHAP-based interpretability tools and interactive 3D feature scatter plots were not used directly for feature selection.

### Class-conditional feature distribution plots

To assess the class separability of each feature in the absence of temporal context, we used DeepTRACE’s *Feature Visualisation* tool, which provided a visual indication of class separability using histograms and kernel density estimates (**Extended Data Fig. 1a**), together with various quantitative metrics described below. Histograms were constructed by adaptive binning of feature distributions into 25 bins with edges computed globally across all tracks and classes to ensure reliable sampling despite extreme differences in feature distributions, scenarios with large class imbalance, and small datasets.

The two-sample Kolmogorov-Smirnov statistic (*D*_KS_) between the empirical cumulative distribution functions of feature values for each class, which can be computed directly on raw feature data without binning, was defined as,

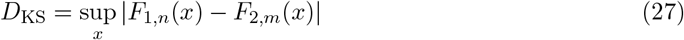

Where *F*_1_ and *F*_2_ represent the cumulative distribution functions of the classes being compared, containing *n* and *m* samples respectively. The 1^st^ order Wasserstein distance (*W*_1_) was computed from adaptively binned data as,

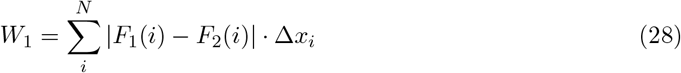

Where *N* is the number of bins, and ∆*x*_*i*_ is the width of the *i*^th^ bin. Finally, the more visually intuitive histogram-based metric (Ω) computed the fractional non-overlap between classes, defined as,

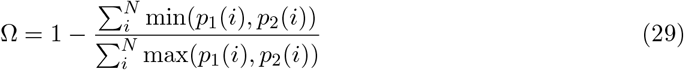

Where *p*_1_(*i*) and *p*_2_(*i*) are occupancies of the *i*^th^ bin in class 1 and 2 respectively.

### Pairwise Pearson Correlation

To identify redundant features for removal, linear correlations were computed using DeepTRACE’s *Feature Ranking* tool. The pairwise Pearson correlation between every feature pair in the dataset was computed across all tracks using,

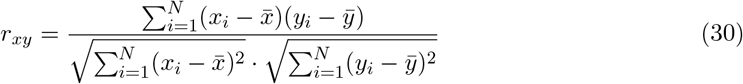

Where *x*_*i*_ and *y*_*i*_ represent the *i*^th^ values of two features, 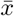 and 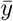 are their mean values, and *N* is the total number of localisations in the combined dataset. The resulting correlation matrix was visualised as a heatmap (**Extended Data Fig. 2a**), enabling identification of strong linear correlations which indicates high redundancy.

### Mutual information

Feature importance was assessed via the mutual information shared by each feature with the class assigned from human annotation or ground truth. To account for the wide diversity of feature distributions, including those with extreme variations in kurtosis (heavy tails or sharp peaks), we discretise the data using adaptive binning into 25 equally-populated bins, such that each distribution is sampled at intervals of even density throughout the dataset. We then compute the mutual information metric as follows,

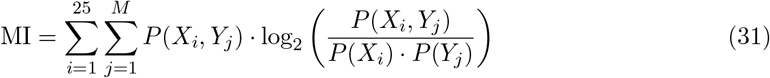

Where *X* represents the discretised feature, *Y* is an integer assigned to each of *M* classes, *P* (*X*_*i*_, *Y*_*j*_) is the joint probability mass function of *X* and *Y*, and *P* (*X*_*i*_) and *P* (*Y*_*j*_) are the marginal probability mass functions of the feature *X* and class *Y*. Pairwise mutual information heatmaps were generated by computing the mutual information between all feature pairs (*X*_*i*_, *Y*_*j*_), where *Y* in this case represents the second feature.

### Training models

#### Preparation of training data

All features were standardised using Z-score normalisation globally across all tracks prior to splitting training, validation, and test data. The Z-score transform rescaled the data to a mean of zero, and standard deviation of one, such that for a given feature *x*, each datapoint *x*_*i*_ was transformed to its Z-score *z*_*i*_ as,

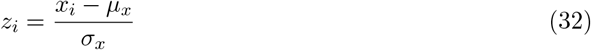

Where *μ*_*x*_ and *σ*_*x*_ represent the global mean and sample standard deviation of the feature x respectively,

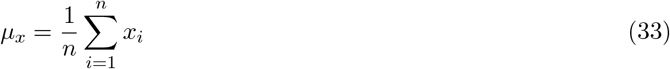

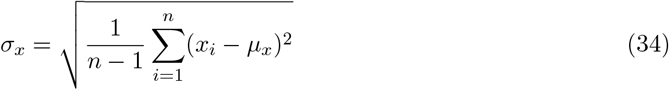

and *n* is the number of timepoints across all tracks in the dataset. While DeepTRACE offers alternative forms of data normalisation, throughout this work we use Z-score as this provides the most robust handling for feature sets with widely differing distributions. These parameters used were stored with the trained model, ensuring consistent preprocessing during evaluation of unseen data.

To enforce sample independence, data shuffling was performed at the individual track level prior to splitting into training, validation, and optional test data. A split of 85% training and 14% validation data was used, and a small test split (1%) was discarded due to the current internal configuration of the training framework; however, all test results reported in this study were obtained from completely separate experimental and simulated datasets, providing a truly independent and more robust assessment of model generalisation than a simple split of the training data. Drawing test data from separate experiments was also necessary as single-molecule tracking data often contain information and artefacts specific to the recording (e.g., environmental conditions, sample heterogeneity, hardware variability, or optical effects such as focal plane positioning).

Given the small dataset sizes, sequences used for training were truncated by two timepoints at both ends to ensure only regions with complete feature coverage were used for training, accelerating the learning process by avoiding learning special cases related to end point proximity.

DeepTRACE offers a range of subsampling methods to compile training data; throughout this work we use the ‘*random subsampling* ‘ method with a temporal context window of 25 localisations. DeepTRACE performed subsampling by computing the maximum number of overlapping subtracks in the shortest track of the dataset, and used this number to randomly subsample without replacement all tracks into the same number of examples. This unusual temporal augmentation strategy increases sample diversity by exposing each datapoint to the largest possible range of local contexts while ensuring that equal weighting was provided to each track regardless of length, maximising the impact of limited data. Resulting examples were then shuffled randomly into batches after separation at the whole-track level of the training and validation data. Note that the concept of training epoch varies slightly with this augmentation approach as unique short sequences appear multiple times in time-shifted contexts across different batches.

#### Model selection

While DeepTRACE supports a range of models, including recurrent neural networks (RNNs) and ensemble-based methods, throughout this study we chose Bidirectional Long Short-Term Memory networks (BiLSTMs) followed by a self-attention mechanism. Throughout early testing this configuration consistently outperformed Gated Recurrent Unit (GRU) and ensemble decision tree models with feature-engineered temporal context.

Additional model types remain available in DeepTRACE for applications beyond the scope of this study, including random forests for analysis of processes lacking temporal dependencies (e.g. fixed cell experiments), and model interpretability tasks.

We conducted empirical benchmarking of different configurations, including models with deeper RNN stacks and multi-head attention mechanisms, but observed no consistent improvement on the small datasets targeted by this study. While the DeepTRACE GUI supports flexible construction of all such models based around a modular architecture (**Extended Data Fig. 10**), all models presented here use a single BiLSTM layer and single attention head. This design prioritises reduced complexity, rapid training on small-scale datasets, and compatibility with limited computing resources, to align with DeepTRACE’s goal of broad accessibility across the single-molecule tracking community.

RNN layers in all trained models presented here used 100 hidden units, a post-RNN dropout layer with 50% dropout, input and recurrent L2 regularisation factors of 2×10^−5^ and no L2 penalty on biases. Initial hyperparameter ranges were explored with a custom-built, Bayesian-inspired search implemented during early development, and final configurations were selected via manual tuning. BiLSTM layers used gating mechanisms (tanh for state updates and sigmoid for gating) that are suited to learning long-range patterns in time series data. Models used in the experimental permutation tests for example (**Extended Data Fig. 9a**) contained 125,202 trainable parameters.

#### The training process

Training used the Adam optimiser with piecewise learning rate scheduling. The initial learning rate was set to 10^−4^ and reduced to 30% of the previous rate every five epochs. All models were trained with mini-batches of 16 examples. Validation loss and accuracy were monitored using a holdout validation set evaluated every 500 iterations. Early stopping was triggered following 10 consecutive evaluations without improvement, or after a maximum of 50 epochs. When working with datasets containing fewer than 250 tracks, and for all models in the experimental perturbation simulations, the validation patience and maximum epochs were extended to 15 consecutive evaluations, and 100 epochs respectively. The final model weights are those that produced the lowest validation loss. Final model evaluation was performed on a completely independent test set from separately-run simulations and experiments.

#### Hardware

Models for the reversible two-state diffusion simulations (**Fig. 2b,c**; **Extended Data Fig. 6c**) were trained using a single core of a typical laptop single CPU (M2 Macbook Air, 2022) in MacOS. All other models were trained using a low-cost GPU (Nvidia Quadro P1000, released February 2017, 4 GB GDDR5 memory, 82 GB/s bandwidth, with no Tensor Cores and no accelerated FP16 support) on a typical Windows desktop PC while running other background tasks. This low-powered, dated GPU was chosen intentionally to demonstrate method efficacy on modest hardware; by today’s standards, this GPU — desgined for lightweight CAD tasks — is outdated and lacks the computational power and memory required for large-scale deep learning workloads.

#### Loss functions

The total loss was computed as the contribution *L*_*b*,*t*_ for all timepoints, *T*, across the mini-batch *B*,

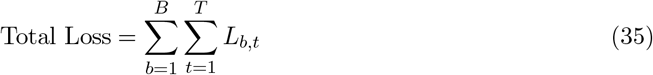

The class-weighted loss was computed using cross-entropy with weights obtained from the inverse of normalised class frequency, to obtain the loss per localisation instance as follows,

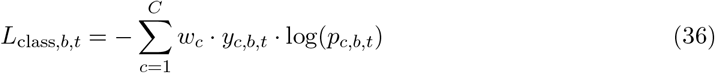

Where *w*_*c*_ is the class weight for class *c*; *y*_*c*,*b*,*t*_ is the true label (1 or 0) for class *c*, one-hot encoded at batch index *b*, and time point *t*; and *p*_*c*,*b*,*t*_ is the predicted probability.

We also defined a changepoint-masked loss function that computes class-weighted cross-entropy loss for a single data point (indexed by *b, t*, and *c*) with an extra weight multiplier for changepoint-proximal regions. Changepoints were defined as any change in ground truth label in adjacent localisations (i.e. *y*_*b*,*t*_ ≠ *y*_*b*,*t*−1_). The combined class-weighted, and changepoint-mask weighted loss contribution for per localisation instance was defined as,

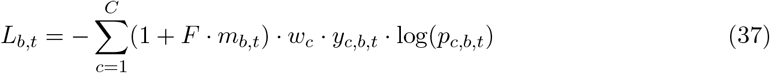

Where *F* is the additional weighting assigned to localisations indexed by the changepoint mask; *m*_*b*,*t*_ is the entry from the binary changepoint mask at batch index *b* and time point *t*. Throughout this work we primarily employ the class-weighted loss as most applications concern the extraction of global biophysical parameters from analysed tracks, which was found to be relatively insensitive to minor increases in changepoint error.

#### Inference

DeepTRACE supports several inference modes, including whole-track and segment-based classification; however, all results presented in this study use its sliding window method. Each track was divided into overlapping temporal windows, and the trained model was applied independently to each window to predict class probabilities using sequence-to-sequence classification. Since localisations appear in multiple overlapping windows, DeepTRACE computes a consensus classification by averaging the softmax probabilities across all windows in which each localisation appears. This allows each annotation to reflect multiple temporally shifted contexts, improving robustness to local noise. The final assigned class corresponds to the class with the highest mean probability.

In addition to the predicted class, DeepTRACE also retains the full probability distribution and number of contributing windows for each localisation, enabling confidence-aware post-processing (e.g. Gaussian or exponential smoothing, enforcing minimum time per state, or enabling state reassignment based on permitted event sequences) if desired. However, to faithfully evaluate the raw performance of each model, no post-processing was applied as this may artificially boost the reported performance, and we instead report on raw performance metrics which reflect the underlying model classification.

#### Visual verification of model performance

To avoid operating as a black box, DeepTRACE’s classification performance was monitored via multiple evaluation metrics, discrete statistics, comparison to human and ground truth data, and visual interrogation of individual tracks. Following segmentation, DeepTRACE was used to perform extensive downstream analysis of segmented tracks, including mobility analysis, spatial mapping, descriptive statistics, and event rates.

#### Performance metrics

Classification performance was quantified using mean changepoint error and various macro-averaged metrics. If *TP*, *FP*, *TN*, and *FN* are the number of true positives, false positives, true negatives, and false negatives respectively, then we compute each metric using standard definitions as follows,

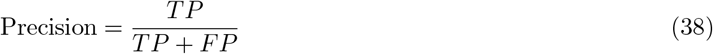

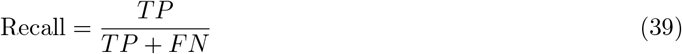

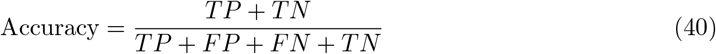

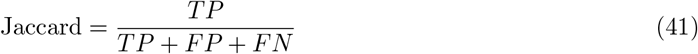

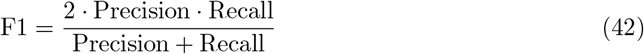

Macro-averaged F1-score was computed as the mean F1-score across all classes. The mean changepoint error was defined as the mean difference in number of frames between matched change-points of the same type in the model classifications and ground truth within a maximum distance of five frames, as follows,

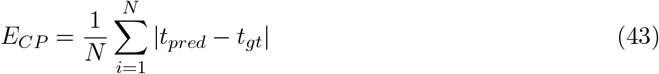

Unmatched changepoints are assigned the maximum changepoint error. Changepoint metrics for real experimental data represent the difference between model annotations and changepoints in the human annotations.

#### Permutation importance

Permutation importance was evaluated using DeepTRACE’s *Permutation Importance* tool, which computed the drop in macro-averaged accuracy of the model across all classes when each feature is permuted. For every feature used by the model, localisation values across all tracks were shuffled, breaking their association with the class labels. Model accuracy was then re-evaluated, and the mean drop in performance across five permutations was recorded along with the standard deviation.

DeepTRACE computed the permutation importance separately within the changepoint-proximal and changepoint-distal regions (defined as the four localisations closest to each ground truth changepoint), and ranked features by the sum of their importance across both regions to identify features most heavily influencing the model. This changepoint masking was particularly instructive as some engineered features were specifically designed for changepoint identification and may not exhibit strong class separation in distal regions, which constitute the majority of frames in most single-molecule tracking data. As with other metrics, ground truth for real experimental data was taken as the human annotations.

#### SHapley Additive exPlanations (SHAP) values

SHAP values (**Extended Data Fig. 2c**) were computed using Random Forest surrogate model with MATLAB’s Tree SHAP algorithm.

#### Spatial mapping

We used DeepTRACE’s *Spatial Mapping* tool to generate 2D heatmaps (**Extended Data Fig. 4c** inset), STORM-like reconstructions (**Extended Data Fig. 3b,c**), and cell-axial projections of classified molecule localisations. Localisations were transformed into model cell coordinates using the *longitude* and *latitude* features engineered by DeepTRACE, with those falling outside the segmented cell boundary excluded from visualisation. Heatmaps were constructed for each class by binning spatial localisations with a bin size of 25 nm. Reconstructed spatial maps were produced by positioning radially symmetric Gaussian PSFs at each localisation within a model cell, and then rendered by accumulating the contribution from all PSFs into a single, normalised super-resolution image with a pixel size equal to half of the PSF’s full width at half maximum (FWHM).

#### Computation of diffusion coefficients

Diffusion coefficients were computed using DeepTRACE’s *Diffusion Analysis* tool, which compiled all segmented subtracks for each class across the entire dataset. Valid displacement steps were aggregated to construct MSD-lag time relationships for each class, and apparent diffusion coefficients were then calculated from a linear least-squares fit to the first two lag times for each class using *D*^∗^ = *MSD/*4*τ*. This approach enables robust diffusion analysis without requiring uninterrupted diffusive states, and greatly reduces the excessive truncation-induced noise that limits conventional single-molecule diffusion analysis. While DeepTRACE is able to compile MSD-lag time relationships by truncating the end points of subtracks to further boost performance (by minimising the influence of class averaging present in localisations adjacent to each end point arising from changepoint errors and intra-frame transitions, see ‘Simulations’ in **Online Methods**), we instead compute coefficients using all available steps to more accurately represent the raw segmented tracks, including these sources of error.

#### Computation of ground truth metrics

Ground truth apparent diffusion coefficients were obtained using standard MSD analysis on complete subtracks segmented using the objective ground truth labels, and computed numerically as a linear fit to lag times of one and two frames as described in ‘Computation of diffusion coefficients’. These values therefore contain the same localisation and statistical noise as is present in the tracking data supplied to Spot-On, γMM analysis, and DeepTRACE. Differences between ground truth apparent diffusion coefficients and the simulation input values are a result of a number of sources of noise, including spatial confinement, statistical noise in drawing from distributions (e.g. state transition probability, initial proximity to spatial barrier, step size), projection effects, and localisation noise; the real-world analogues of each exist in experimental data.

#### Evaluation with Spot-On

To obtain a fair comparison to the widely-used software Spot-On, we analysed the exact set of filtered tracks used by DeepTRACE, reformatted to the Spot-On default format (*Frame, Time, Track ID, x, y*). Spot-On’s two-state model was used for all analysis, with default parameters, which were empirically well-suited to our simulation data. The ranges of *D*_bound_ and *D*_free_ were adjusted based on known ground truth values, and bin widths were tuned to improve sampling resolution. Fitting was also performed with both CDF and PDF methods, and using truncated and non-truncated forms to ensure the best possible configuration was obtained.

#### Gamma Mixture Model analysis using truncated tracks

Tracks with a minimum length of five localisations were used to compile diffusion histograms, with longer tracks truncated to this minimum length. For each track, the apparent diffusion coefficient *D*^∗^ was estimated by performing a linear fit to the MSD-lag time relationship and aggregated to populate diffusion histograms. A two-Gamma mixture model was fit to the ensemble diffusion histogram to estimate the underlying distribution of apparent diffusion coefficients. Following the method of Stracy et al. [15], we fit the probability density function 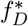 of two Gamma distributions with diffusion coefficients *D*_1_ and *D*_2_, and occupancies *A*_1_ and *A*_2_ respectively,

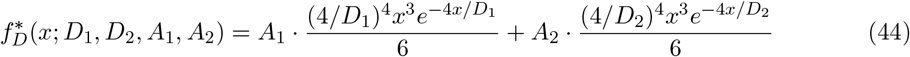

We leave both diffusion coefficients and occupancies as free parameters to avoid biasing the fitting process. To retain consistency with previously published methods, Gamma mixture model analysis was performed on the raw tracking data; as such the analysis contains additional short tracks that were removed by DeepTRACE’s track filtering processes.

#### Construction of diffusion histograms from DeepTRACE-segmented subtracks

Diffusion histograms were constructed with DeepTRACE’s *Diffusion Histogram* tool, which aggregates data from subtracks grouped by class. Each histogram entry corresponds to the apparent diffusion coefficient of a single subtrack computed via linear least squares regression on the first two entries of the mean MSD-lag time relationship, using all available steps within the subtrack.

While all diffusion coefficients presented were computed directly from a single fit to an aggregated MSD-lag time plot for each class (see ‘Computation of diffusion coefficients’), coefficients can also be estimated, albeit less accurately, by fitting Gaussian components to each segmented diffusion histogram (**Extended Data Fig. 6b**).

## Acknowledgements

We thank Valentine Lagage and Stephan Uphoff for providing access to the published DNA polymerase I experimental dataset. We are grateful to Alison Farrar and Mirjam Kummerlin for providing independent human annotations during the development of DeepTRACE. We also thank Mathew Stracy, Isabelle Lorge, Stephan Uphoff, and Sebastian von Hausegger for their feedback on the manuscript.

## Author contributions

O.J.P. conceived the DeepTRACE concept and programmed the DeepTRACE software. O.J.P. designed and trained all models. O.J.P. performed the data analysis. The simulation pipeline was co-designed by O.J.P. and J.A.R.W.. J.A.R.W. programmed the simulation pipeline, and generated all simulated data. The manuscript and figures were written by O.J.P. with feedback and revisions from A.N.K. The Online Methods section covering simulations was written by J.A.R.W. with revisions by O.J.P.. The project was supervised by A.N.K. All authors approved the manuscript and its submission.

